# Unveiling the physiological mechanisms that drive the emergence of antibiotic resisters from antibiotic persister population of mycobacteria

**DOI:** 10.1101/846378

**Authors:** Kishor Jakkala, Deepti Sharan, Rashmi Ravindran Nair, Avraneel Paul, Atul Pradhan, Parthasarathi Ajitkumar

## Abstract

The physiological mechanisms behind the emergence of antibiotic-resistant bacteria from their antibiotic-persister population are beginning to be explored. Here we delineate the sequential physiological events that drive the emergence of rifampin-resistant *rpoB* mutants from rifampin-persister population of mycobacteria during prolonged exposure. The rifampin-persister population generated elevated levels of hydroxyl radical, which inflicted mutations, enabling regrowth of the persister cells to form multi-septated, multi-nucleated elongated cells. These cells, through multiple divisions, produced multiple sister-daughter cells, causing an abrupt, unexpectedly high increase of rifampin-resistant colonies. Similar response was observed against moxifloxacin also. Earlier studies on the rifampin/moxifloxacin-exposed laboratory/clinical *M. tuberculosis* strains from *in-vitro* cultures and infected mouse-lung also showed cfu spurt, but went unnoticed/unreported. It is likely that these sequential physiological events may be driving the emergence of antibiotic-resistant tubercle bacilli in TB patients also. *Escherichia coli* also has been found to respond similarly against subminimal inhibitory concentrations of ciprofloxacin. Thus, the present findings attain broad significance as a general physiological mechanism used by diverse bacterial genera to emerge as drug-resistant strains against antibiotics.

## Introduction

Antibiotic bacterial persisters were first observed in the recurrence of Staphylococcal infections, despite extensive treatments with high doses of penicillin (Hobby *et al*., 1942; Bigger, 1944). The persisters constitute a subpopulation of cells in a bacterial population that can tolerate lethal concentrations of antibiotics and, upon removal of the antibiotic, can re-grow to generate back the antibiotic sensitive population. The difference between bacterial persistence and antibiotic resistance is that unlike the antibiotic resistant mutants, which grow in the presence of the antibiotic/bactericidal agent, the persister cells do not proliferate in the presence of the antibiotic/bactericidal agent, but resume growth once the antibiotic is removed, giving rise to antibiotic-sensitive population again (Bigger, 1944). The antibiotic-sensitive population can once again generate persisters in the presence of the antibiotic. Thus, persistence, in contrast to resistance, is a non-inherited phenomenon.

Bacteria, both pathogenic and non-pathogenic, of diverse genera and habitat have shown the phenomenon of persistence. With mycobacteria being no exception to this trait, large number of studies have been carried out on *Mycobacterium tuberculosis* persisters (Robertson, 1933; McCune *et al*., 1966a, b; Hu *et al*., 2000; Lenaerts *et al*., 2007; Hoff *et al*., 2011; Sebastian *et al*., 2017). High density of acid-fast bacilli has been found in the tubercles resected months after the patients on chemotherapy had become sputum negative (Medlar *et al*., 1952; Beck and Yegian, 1952). Similarly, tubercle bacilli have been found in the lung tissues taken at autopsy from individuals who died from reasons unrelated to TB (Opie and Aronson, 1927; Robertson, 1933; reviewed in Stewart *et al*., 2003). Ovoid shaped, “ultra-fine” forms of infectious bacilli, revivable under certain growth conditions, have been found in the sputum and plasma of TB patients receiving anti-TB therapy (Khomenko, 1987). Mice infected with virulent *M. tuberculosis* H37Rv contained *in vitro* drug-susceptible persistent bacilli in the lungs and spleen despite treatment with anti-TB drugs (McCune and Tompsett, 1956a; McCune *et al*., 1956; McCune *et al*., 1966a; McCune *et al*., 1966b). The bacterial load in the mouse model was found to remain the same during the persistence of the bacilli indicating non-replicative nature of the persister cells (Rees and D’Arcy Hart, 1961). *M. tuberculosis* infected guinea pig contained persistent bacilli in the lungs, spleen and in the extracellular milieu, regardless of drug treatment (Lenaerts *et al*., 2007; Kashino *et al*., 2008; Hoff *et al*., 2011). Persistent *M. tuberculosis* has also been observed in infected macrophages (McKinney *et al*., 2000; Adams *et al*., 2011) and in i*n vitro* cultures (Hu *et al*., 2000; Singh *et al*., 2010; Keren *et al*., 2004; Sebastian *et al*., 2017).

There are many studies that indicate possible association of persistence with drug resistance. Accumulation of point mutations associated with antibiotic resistance developed over several months during treatment has been demonstrated using whole-genome sequencing of *Staphylococcus aureus* isolates from a patient with persistent infection (Mwangi *et al*., 2007). Similarly, whole-genome sequencing of *M. tuberculosis* populations in the sputum from patients, who experienced recrudescent infection, revealed hetero-resistance of *M. tuberculosis* population with newly acquired resistance (Sun *et al*., 2012). Although multiple resistant mutants transiently coexist within the population, only a single type of mutant that is highly adaptive and retained fitness despite the cost towards incurring mutations would get selected and survive (Sun *et al*., 2012). According to the recent WHO report on TB, 20% of the retreatment cases harbor MDR-TB, in contrast to 3.3% of new cases, indicating the possible emergence of resistant mutants in the retreatment cases from uncleared persisters (WHO Global TB Report, 2015).

It has been proposed that the antibiotic persister cells could behave as an evolutionary reservoir for the emergence of antibiotic-resistant mutants (Cohen *et al*., 2013; Sebastian *et al*., 2017). This has been experimentally demonstrated in diverse bacterial systems (reviewed in Cohen *et al*., 2013), including *M. tuberculosis* and *M. smegmatis* (Cohen *et al*., 2013; Sebastian *et al*., 2017; Swaminath, 2017). The emergence of *M. smegmatis* persisters *in vitro* upon treatment with rifampin, INH, ciprofloxacin, and ofloxacin has been reported (Grant *et al*., 2012; Swaminath, 2017). We have recently shown the *de novo* emergence of rifampin/moxifloxacin-resistant genetic mutants of *M. tuberculosis* and *M. smegmatis* from the respective antibiotic persistence phase cells due to mutagenesis from elevated levels of hydroxyl radical (Sebastian *et al*., 2017; Swaminath, 2017). The genetically resistant mutants emerged from the persister population of cells despite the continuous presence of microbicidal concentrations of the antibiotic for prolonged duration, such as for 20 days for *M. tuberculosis* and 96 hrs for *M. smegmatis* cells *in vitro* (Sebastian *et al*., 2017; Swaminath, 2017). The antibiotic-resistant genetic mutants regrew and populated the culture in the continued presence of microbicidal concentrations of the antibiotic.

In the background of these studies, we addressed here as to what are the sequential physiologic events that propel the persister cells of mycobacteria against rifampin and moxifloxacin to acquire mutations against antibiotics, regrow, divide, and generate a population of antibiotic-resistant genetic mutants in the continuous presence of the respective antibiotic. Since identical response in terms of sudden spurt in cfu was shown by the antibiotic-resistant genetic mutants of both *M. smegmatis* (saprophyte) and *M. tuberculosis* (avirulent H37Ra and virulent H37Rv) also, we discussed the relevance of this finding in the emergence of drug-resistant tubercle bacilli in TB patients.

## Results

### Background information and experimental strategy

We had earlier found that *M. tuberculosis* (*Mtb*) and *M. smegmatis* (*Msm*) cells, which were exposed to minimum bactericidal concentrations (MBCs) of rifampicin and moxifloxacin for prolonged duration *in vitro* such as 18 days for *Mtb* cells and 96 hrs for *Msm* cells, showed a sequential killing, persistence and regrowth phases (Sebastian et al, 2017; Swaminath, 2017). In these studies, rifampicin-resistant and moxifloxacin-resistant genetic mutants of *Mtb* and *Msm* were found to emerge from the persister population, regrow and populate the culture. The rifampicin-resistant and moxifloxacin-resistant mutants were having nucleotide changes at the rifampicin resistance determining region (RRDR) and quinolone resistance determining region (QRDR), respectively (Sebastian et al, 2017; Swaminath, 2017). These mutations were identical to and at identical positions as those observed in the rifampicin-resistant and moxifloxacin-resistant clinical isolates of *Mtb* (Cavusoglu *et al*., 2002; Takiff *et al*., 1994). Since this phenomenon was found in both *Mtb* and *Msm* cells and irrespective of the antibiotic used, the present study was performed using *Msm* cells only for faster pace of getting colonies and response from the cells against antibiotics. However, to present the response of *Mtb* cells to rifampicin and moxifloxacin, we have considered the cfu values corresponding to the graphs depicting the *Mtb* susceptibility to rifampicin and moxifloxacin, respectively, from the earlier published work from our laboratory (Sebastian et al, 2017; Copyright @ American Society for Microbiology, Antimicrobial Agents and Chemotherapy, 61, 2017, e01343-16, https://doi.org/10.1128/AAC.01343-16). In this work, the unexpectedly high levels of cfu spurts that occurred against rifampicin and moxifloxacin were not reported as they went unnoticed by us.

Thus, as per the rationale mentioned above, we exposed *Msm* mid-log phase (MLP) cells individually to MBCs of rifampin or moxifloxacin and plated on the antibiotic-free and/or antibiotic-containing (125 µg/ml rifampin; 3x MBC, Sharmada, 2017) Mycobacteria 7H11 agar plates once every 6 hrs for 120 hrs for the determination of total cfu and the cfu of antibiotic-resistant mutants, respectively. Depending upon the conspicuous changes in the cfu on the antibiotic-free plates during the entire exposure period, the killing phase with exponential reduction in the cfu, the persistence phase with no appreciable change in the cfu, and the regrowth phase showing steady rise in the cfu, were temporally demarcated. The different phases were identified to isolate the cells from the respective phase for the analyses.

The colonies from the rifampin-free plates were patch-plated onto rifampin-containing plates to identify which of the colonies that grew on rifampin-free plates were actually rifampin-resistant. These colonies from both the antibiotic-free and antibiotic-containing plates were selected from five time points for RRDR sequence determination. These time points included the time point showing abrupt several-fold high spurt in cfu, two time points prior to and two time points later to the time point of sudden spurt in cfu. The cells from these time points were examined using transmission electron microscopy, atomic force microscopy, fluorescence microscopy, and live-cell imaging for cell division, filamentation, if any, and for the presence of multiple nucleoids and multiple septae. The number of cells with ≤2n and >2n nucleoids were quantitated for the cells from these five time points. The initial cell density chosen for the experiment was 10^6^ cells/ml to pre-empt the natural resisters to rifampicin that exist at 10^−8^ frequency in the MLP culture (Swaminath, 2017). All the experiments were performed using biological triplicate samples and statistical significance of the observations was calculated. Similar analyses were performed for the *Msm* cells exposed to 3.75x MBC moxifloxacin for prolonged duration, as reported (Swaminath, 2017). However, the mutations in the *gyrA* gene in the moxifloxacin-resistant mutants formed from the persisters were not determined since the response of the bacilli to moxifloxacin was identical to that to rifampin. Further, the mutations in the moxifloxacin-resistant genetic mutants formed from the persisters were already determined in the recent study on *Msm* persisters against moxifloxacin from our laboratory (Swaminath, 2017).

### The response of *Msm* cells to MBCs of rifampin and moxifloxacin

The rifampin susceptibility curve obtained from the cfu on the rifampin-free plate showed a killing phase, with a steady decline in the cfu (∼3-log10 reduction in the cfu), from 0 hr to 36 hr (**Fig. 1A**). The killing phase was followed by the persistence phase, from 36 hr to 54 hr, characterised by practically no appreciable change in the cfu (**Fig. 1A**). The persistence phase was ensued by the regrowth phase, from 54 to 120 hr, where a steady rise in the cfu by ∼4-log10 difference could be observed (**Fig. 1A**). The temporal demarcation of the three phases in terms of the change/no change in the cfu enabled isolation of the cells from the persistence and regrowth phases for further analyses. A response, similar to that shown to rifampin, was found from the *Msm* cells to MBC of moxifloxacin (0.5 µg ml^−1^; 3.75x MBC) upon prolonged exposure for 96 hrs (**Fig. 1B**), as reported in our recent study (Swaminath, 2017). Here also, in terms of the changes in the cfu from the moxifloxacin-free plates, the killing (from 0 hr to 36 hr), persistence (from 36 hr to 54 hr), and regrowth (from 54 hr to 96 hr) phases could be identified. Thus, despite the modes and targets of action of rifampin and moxifloxacin being different, *Msm* cells showed identical response to them, with the emergence of regrowing population in the continued presence of MBCs of the respective antibiotics.

**Figure 1.**
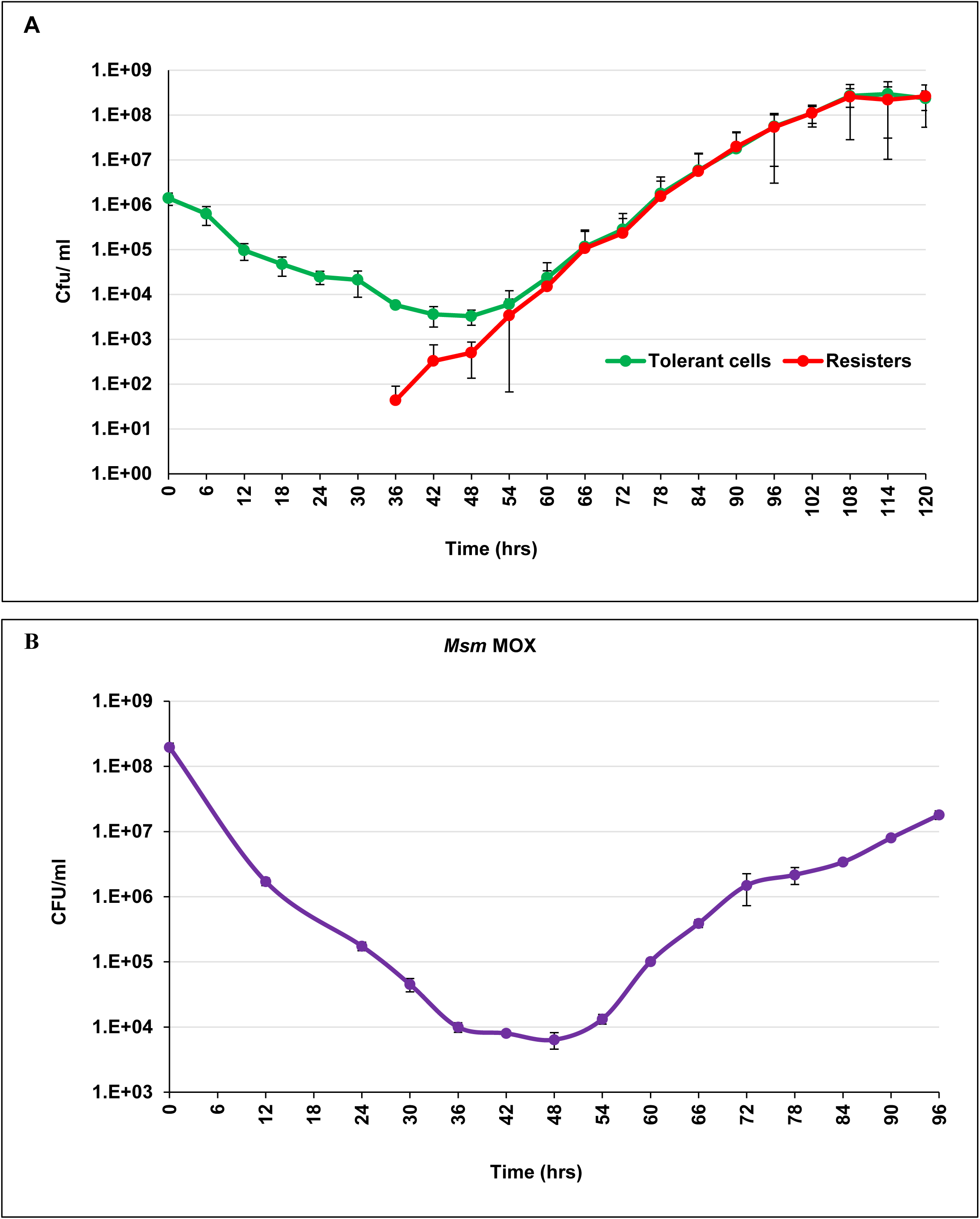
Response of *Mycobacterium smegmatis* cells to rifampin and moxifloxacin upon prolonged exposure. Response to: **(A)** rifampin (25 µg ml^−1^) and **(B)** moxifloxacin (0.5 µg ml^−1^). The profiles show clear killing phase (0 hr to 36 hr with steep decrease in the cfu), followed by the persistence phase (36 hr to 54 hr with no appreciable change in the cfu), and continued by the regrowth phase (from 54 hr till 96/120 hr with steady rise in cfu). The red line in **(A)** indicate the cfu of rifampin-resisters that emerged from the persister population. The moxifloxacin-resisters were not scored (see text for explanation).

### The response of *Mtb* cells to MBCs of rifampin and moxifloxacin

Earlier, we had shown the response of *Mtb* cells upon prolonged exposure (20 days) to microbicidal concentrations of rifampin (10x MBC) and moxifloxacin (2x MBC) (**Fig. S1A,B** here, which are Fig. 1A and Fig. S2 in Sebastian et al, 2017, taken with copyright permission from AAC). The susceptibility profile of *Mtb* cells to MBCs of rifampicin and moxifloxacin also showed killing, persistence and regrowth phases, wherein the regrowing cells were found to emerge from the persister population, like in the case of *Msm* cells exposed to the two antibiotics. Further, our recent studies on *Mtb* and *Msm* cells exposed to rifampicin and moxifloxacin for prolonged duration showed that the cells regrowing from the respective persister population in the continued presence of MBCs of the antibiotics were antibiotic-resistant genetic mutants carrying mutations in the *rpoB* and *gyrA* targets (Sebastian et al, 2017; Swaminath, 2017).

This dynamic change in the respective antibiotic persister population from a non-growing, non-dividing, but antibiotic-tolerant state to an actively growing and dividing antibiotic-resistant state offered an excellent experimental system to identify and characterise the physiological events that accompanied such dynamic change that led to the emergence of antibiotic-resistant mutants. Further, the comparable response of *Msm* and *Mtb* cells to the same antibiotics indicated that the nature of the response of mycobacteria upon prolonged exposure to microbicidal concentrations of antibiotics is an inherent trait that is not influenced by the species type, saprophytic or pathogenic character of the bacterium or by the nature of the antibiotics or the differences in the targets or the cellular processes affected by them. Moreover, such similarity also indicated commonality in the mechanisms involved, justifying the use of *Msm* as the model system to study the physiological events that drive the emergence of antibiotic-resistant mutants from the persister population.

### Abrupt abnormally high spurt in the cfu of antibiotic-tolerant *Msm* cells regrowing from the persister population

The cfu of the persister phase cells (36 hr to 54 hr) on the rifampin-free plates did not show any notable increase (**Fig. 2A, n = 3; Table 1A, rifampin replicates**). The cfu started increasing from 60 hr, with the value abruptly spurting 2-3-fold above the expected cell number doubling consistently during 90-96 hr period (**Fig. 2A, n = 3; Table 1A, rifampin replicates**). Since the cell number doubling time of *Msm* cells in nutrient broth is ∼3 hrs (Gadagkar & Gopinathan, 1980), a 2-fold increase in the cfu was expected in 3 hrs and 4-fold in 6 hrs. But unexpected, unusually high, 2-3 times higher than the 4-fold (i.e., 8-10-fold higher), increase in the cfu was observed in 6 hrs in the rifampin-tolerant cells (**Fig. 2A, Table 1A; n = 3**).

**Figure 2.**
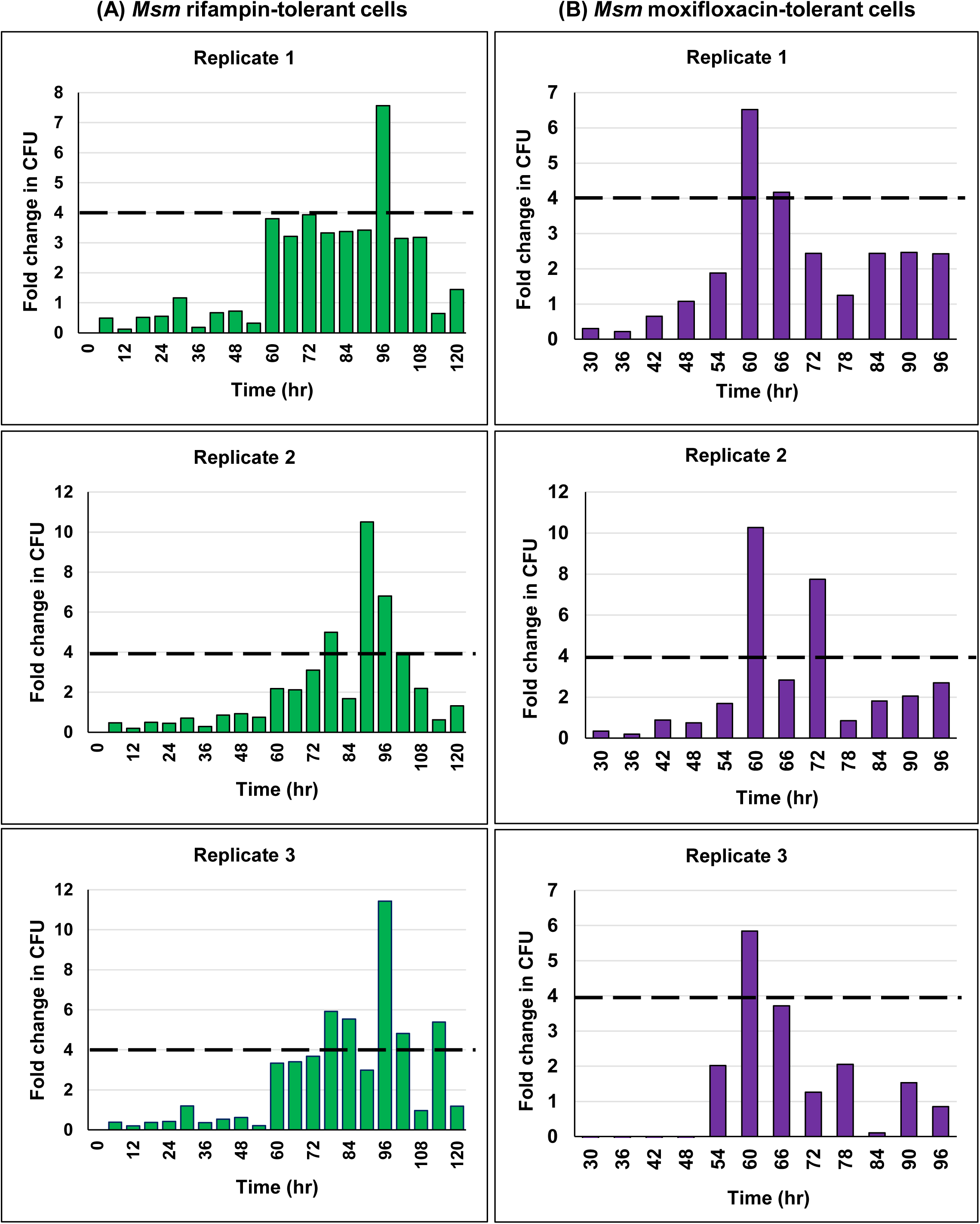
Response of *M. smegmatis* upon prolonged exposure to rifampin and moxifloxacin. Rifampin and moxifloxacin were used at 25 µg ml^−1^ and 0.5 µg ml^−1^, respectively, for exposure. The cfu, from rifampin-free and moxifloxacin-free Mycobacteria 7H11 agar plates, every 6 hrs during the exposure, was determined. The fold-increase in the cfu, with respect to the cfu of the previous time point, was plotted for: **(A)** rifampin-exposed and **(B)** moxifloxacin-exposed cells. The dotted line indicates the maximum expected increase in the cfu in 6 hr period (i.e., 4-fold, as the generation time of *Msm* cells is 3 hrs). n = 3 biological replicates in both cases.

**Table 1.** CFU values of *M. smegmatis* cells exposed to MBCs of rifampin and moxifloxacin. Tables showing the observed and the expected CFU values (in parenthesis) of 3 biological replicates of *Msm* cells exposed to MBCs of rifampin and moxifloxacin at every 6-hr time interval on the respective antibiotic-free Mycobacteria 7H11 agar plates for three biological replicates. Considering that the maximum expected fold-increase in the CFU of *Msm* cells will be 4-fold in 6 hrs (generation being 3 hrs), the time points showing an abrupt increase in the CFU, which is more than the expected value (from two doublings in cell number) are highlighted in red coloured boxes. Except for the 90-96 hrs period for the rifampin-exposed cells and for the 54-60 hrs period for the moxifloxacin-exposed cells, the time points for the spurt in CFU varied among the replicates. The 0-24 hr period for moxifloxacin-exposed samples are not shown as there was no increase in cfu (see Fig. 2B).

| Rifampin replicate 1 |  | Rifampin replicate 2 |  | Rifampin replicate 3 |  |
| --- | --- | --- | --- | --- | --- |
| Time (hrs) | Observed CFU | Time (hrs) | Observed CFU | Time (hrs) | Observed CFU |
| 0 | 1.52E+06 | 0 | 1.80E+06 | 0 | 1.72E+06 |
| 6 | 7.10E+05 | 6 | 8.90E+05 | 6 | 6.60E+05 |
| 12 | 1.35E+05 | 12 | 1.05E+05 | 12 | 1.29E+05 |
| 18 | 6.70E+04 | 18 | 5.40E+04 | 18 | 4.80E+04 |
| 24 | 3.00E+04 | 24 | 3.00E+04 | 24 | 2.00E+04 |
| 30 | 2.10E+04 | 30 | 3.50E+04 | 30 | 2.40E+04 |
| 36 | 6.00E+03 | 36 | 6.40E+03 | 36 | 8.80E+03 |
| 42 | 5.10E+03 | 42 | 4.30E+03 | 42 | 4.70E+03 |
| 48 | 4.70E+03 | 48 | 3.10E+03 | 48 | 2.90E+03 |
| 54 | 3.50E+03 | 54 | 1.00E+03 | 54 | 6.00E+02 |
| 60 | 7.60E+03 | 60 | 3.80E+03 | 60 | 2.00E+03 |
| 66 | 1.61E+04 | 66 | 1.22E+04 | 66 | 6.80E+03 |
| 72 | 5.00E+04 | 72 | 4.80E+04 | 72 | 2.50E+04 |
| 78 | 2.50E+05<br>(2.00E+05 was the cfu expected in 6 hrs) | 78 | 1.60E+05 | 78 | 1.48E+05<br>(1.00E+05 was the cfu expected in 6 hrs) |
| 84 | 4.20E+05 | 84 | 5.40E+05 | 84 | 8.20E+05<br>(5.92E+05 was the cfu expected in 6 hrs) |
| 90 | 4.41E+06<br>(1.68E+06 was the cfu expected in 6 hrs) | 90 | 1.85E+06 | 90 | 2.45E+06 |
| 96 | 3.00E+07<br>(1.76E+07 was the cfu expected in 6 hrs) | 96 | 1.40E+07<br>(7.40E+06 was the cfu expected in 6 hrs) | 96 | 2.80E+07<br>(9.80E+06 was the cfu expected in 6 hrs) |
| 102 | 1.19E+08 | 102 | 4.40E+07 | 102 | 1.35E+08 |
| 108 | 2.60E+08 | 108 | 1.40E+08 | 108 | 1.30E+08 |
| 114 | 1.60E+08 | 114 | 9.00E+07 | 114 | 7.00E+08<br>(5.20E+08 was the cfu expected in 6 hrs) |
| 120 | 2.10E+08 | 120 | 1.30E+08 | 120 | 8.30E+08 |

**Table 1. CFU values of *M. smegmatis* cells exposed to MBCs of rifampin and moxifloxacin.**
| Moxifloxacin replicate 1 |  | Moxifloxacin replicate 2 |  | Moxifloxacin replicate 3 |  |
| --- | --- | --- | --- | --- | --- |
| Time (hrs) | Observed CFU | Time (hr) | Observed CFU | Time (hr) | Observed CFU |
| 30 | 5.43E+04 | 30 | 4.89E+04 | 30 | 3.21E+04 |
| 36 | 1.19E+04 | 36 | 9.80E+03 | 36 | 8.20E+03 |
| 42 | 7.80E+03 | 42 | 8.70E+03 | 42 | 7.50E+03 |
| 48 | 8.40E+03 | 48 | 6.50E+03 | 48 | 4.30E+03 |
| 54 | 1.60E+04 | 54 | 1.10E+04 | 54 | 1.30E+04 |
| 60 | 1.02E+05<br>(6.40E+04 was the cfu expected in 6 hrs) | 60 | 1.13E+05<br>(4.40E+04 was the cfu expected in 6 hrs) | 60 | 8.90E+04<br>(5.20E+04 was the cfu expected in 6 hrs) |
| 66 | 4.30E+05 | 66 | 3.20E+05 | 66 | 4.20E+05 |
| 72 | 1.05E+06 | 72 | 2.48E+06<br>(1.28E+06 was the cfu expected in 6 hrs) | 72 | 9.50E+05 |
| 78 | 1.50E+06 | 78 | 2.10E+06 | 78 | 2.90E+06 |
| 84 | 3.20E+06 | 84 | 3.80E+06 | 84 | 3.20E+06 |
| 90 | 8.00E+06 | 90 | 7.80E+06 | 90 | 8.10E+06 |
| 96 | 1.80E+07 | 96 | 2.10E+07 | 96 | 1.50E+07 |

Although we have observed the unusually high abrupt spurt in the cfu consistently several times (n > 3) during the 90-96 hr period, the abrupt unexpectedly high spurt in the cfu occurred at other periods (72-78 hr, 78-84 hr, 84-90 hr, 96-102 hr, and 108-114 hr) as well, among the replicates (**Fig. 2A, n = 3;** Table 1A**, rifampin replicates**). Hence an averaging of the cfu values for the time points amongst the biological replicates was not attempted. Nevertheless, it was of interest to note that the spurt in the cfu always occurred during the regrowth phase of the persister population, albeit during different periods, but more often during the 90-96 hr period. Essentially, these could be the rifampin-tolerant cells that were regrowing from the persister population, probably after gaining antibiotic-target specific mutation. Since the spurts in the cfu were found at different periods during the regrowth phase, the cfu were formed through fresh rounds of growth and division and not just a carryover of the same number of mutants from one period to the other.

Identical response was shown by the *Msm* cells exposed for prolonged duration to moxifloxacin also. In the moxifloxacin-exposed cells, the cfu spurt to abnormal levels (8-10-fold higher than expected) occurred more often during the 54-60 hr period, although such spurts in the cfu occurred during the 66-72 hr period as well, among the three biological replicates (**Fig. 2B, n = 3;** Table 1B**, moxifloxacin replicates**). Like in the case of rifampin-exposed cells, these cells might be the moxifloxacin-tolerant cells that were regrowing from the persister population, probably after gaining moxifloxacin-target specific mutation.

### Abrupt unusually high spurt in the cfu of rifampin-resistant *Msm* cells regrowing from the persister population

In parallel to plating on rifampin-free plates mentioned above, we had plated an aliquot each of the samples from the same time points on rifampin-containing plates also. Like in the case of rifampin-tolerant cells, the cfu on the rifampin-containing plates also showed a sudden 2-fold to 3-fold higher value (i.e., 8-10-fold higher) than the 4-fold increase in the cfu that was expected in the resisters at different time points, amongst the biological triplicate samples (**Fig. 3A, B; n = 3**). The sudden unexpectedly high spurt in the cfu was observed most often during the 54-60 hr period, although such spurts in the cfu were found during other periods (42-48 hr, 48-54 hr, 60-66 hr, 66-72 hr, 72-78 hr, 78-84 hr, 84-90 hr, 90-96 hr, 96-102 hr, and 108-114 hr) as well (**Fig. 3B; n = 3**). Again, since the cfu spurts occurred at different periods during the regrowth phase, the cfu were contributed by the cells formed through fresh rounds of growth and division. Thus, they were not constituted by a carryover of the same number of mutants from one period to the other. This indicated active growth and division of rifampin-resistant mutants in the continued presence of microbicidal concentrations of the antibiotic.

**Figure 3.**
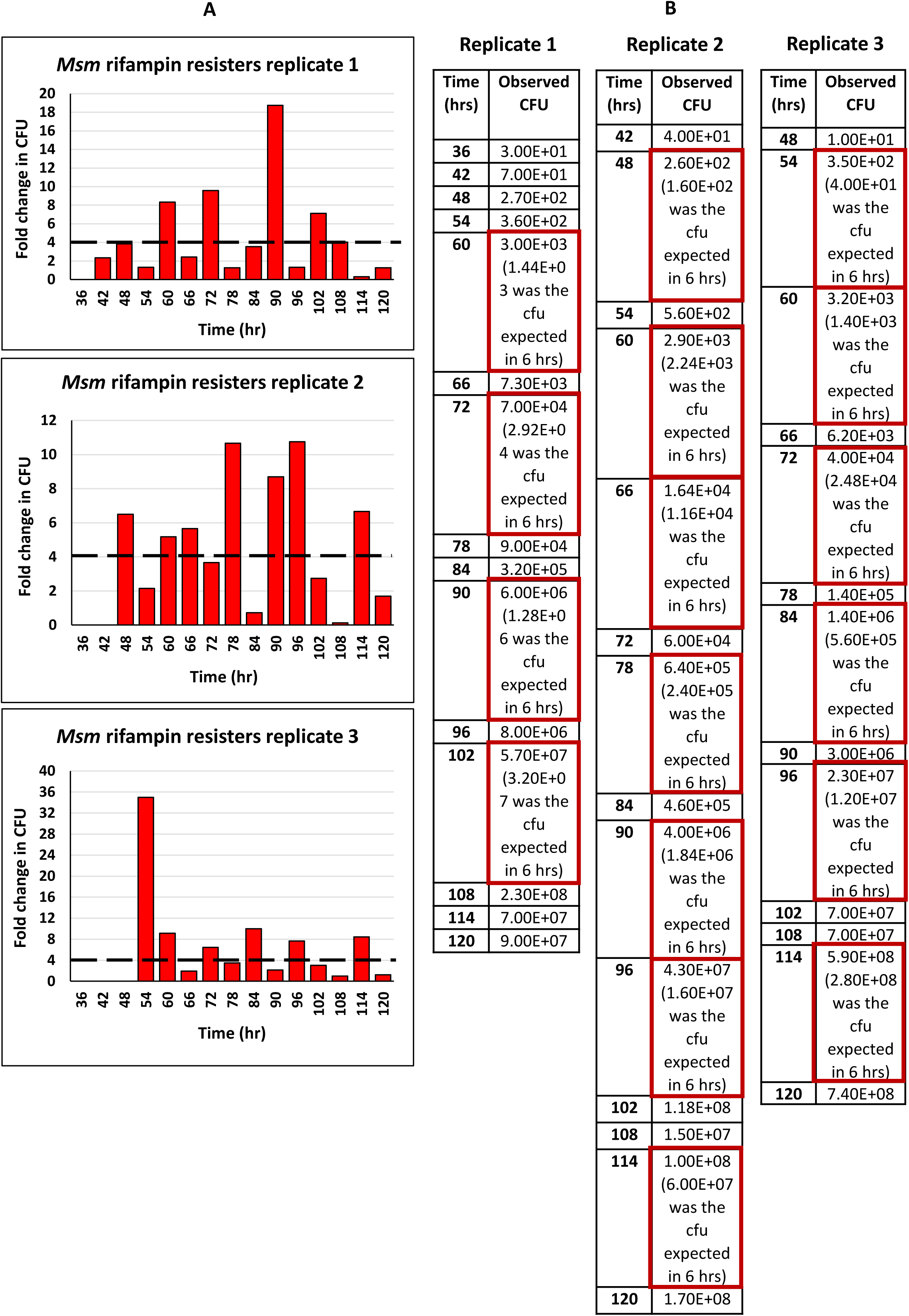
Unusually high level of emergence of rifampin resisters from the cells regrowing from *M. smegmatis* persister population during prolonged exposure to rifampin. CFU and fold-change in CFU of the *Msm* cells exposed to rifampin (25 µg ml^−1^) and plated on rifampin-containing (125 µg ml^−1^) Mycobacteria 7H11 agar every 6 hrs during the exposure. **(A)** The fold-change in CFU compared to the CFU of the previous time point of the biological triplicates. Maximum expected increase in the CFU within 6 hrs (4-fold) is indicated by the dotted line. **(B)** The actual CFU values of the biological triplicates.

### Sudden abnormally high spurt in the cfu of antibiotic-tolerant/resistant *Mtb* cells regrowing from the persister population

We had earlier shown the *de novo* emergence of rifampicin-resistant and moxifloxacin-resistant *Mtb* cells regrowing from the respective persister population of *Mtb* cells formed upon exposure to the respective antibiotic for prolonger duration (Sebastian *et al*., 2017). The persistence phase of the *Mtb* cells exposed to rifampicin and moxifloxacin was found to be from day 10 to day 15 and day 16, respectively, subsequent to which regrowth started (Fig. 1A and Fig. S2 in Sebastian *et al*., 2017; reproduced here as **Fig. S1**, as per the copyright policy of AAC). However, in that work, the spurt in the cfu went unnoticed and hence unreported in the paper (Sebastian *et al*., 2017). When we saw the abnormally high spurt in the cfu of the *Msm* cells regrowing from the persister population in the presence of MBC of rifampin and moxifloxacin, we rechecked the *Mtb* data for the Fig. 1A (for rifampin-tolerant/resistant cells) and Fig. S2 (for the moxifloxacin-tolerant/resistant cells) in the paper (Sebastian *et al*., 2017). Similar to the response of the *Msm* cells exposed to rifampin and moxifloxacin, the *Mtb* cells regrowing from the respective persister population, in the presence of MBCs of rifampin and moxifloxacin, also showed unusually high levels of spurt in cfu (**Fig. 4A, B and C, D, respectively**). The cfu values, pertaining to spurts only, against rifampicin and moxifloxacin, shown here in **Fig. 4**, were taken from the susceptibility profiles of the *Mtb* cells to rifampicin and moxifloxacin, respectively, depicted in **Fig. S1** [taken from the Fig. 1A (for rifampin-tolerant/resistant cells) and Fig. S2A (for moxifloxacin-tolerant/resistant cells) of the paper, Sebastian *et al*., 2017; reproduced here as **Fig. S1**, as per the copyright policy of AAC)]. Thus, the abnormally high spurt in the *Mtb* cfu values of the rifampicin-tolerant/resister cells and moxifloxacin-tolerant/resister cells were found to be from 3-fold to 20-fold (**Fig. 4A, B and C, D, respectively**). Like in the case of the emergence of rifampin/moxifloxacin-tolerant/resister *Msm* cells, the emergence of *Mtb* rifampin/moxifloxacin-tolerant/resister cells also occurred on different days (**Fig. 4A, B and C, D, respectively**). These observations indicated that the unexpectedly high-fold in the cfu of the *Msm* and *Mtb* cells would not have happened but for some unusual mode of cell growth and division occurring during the regrowth phase of the persister cells during antibiotic exposure.

**Figure 4.**
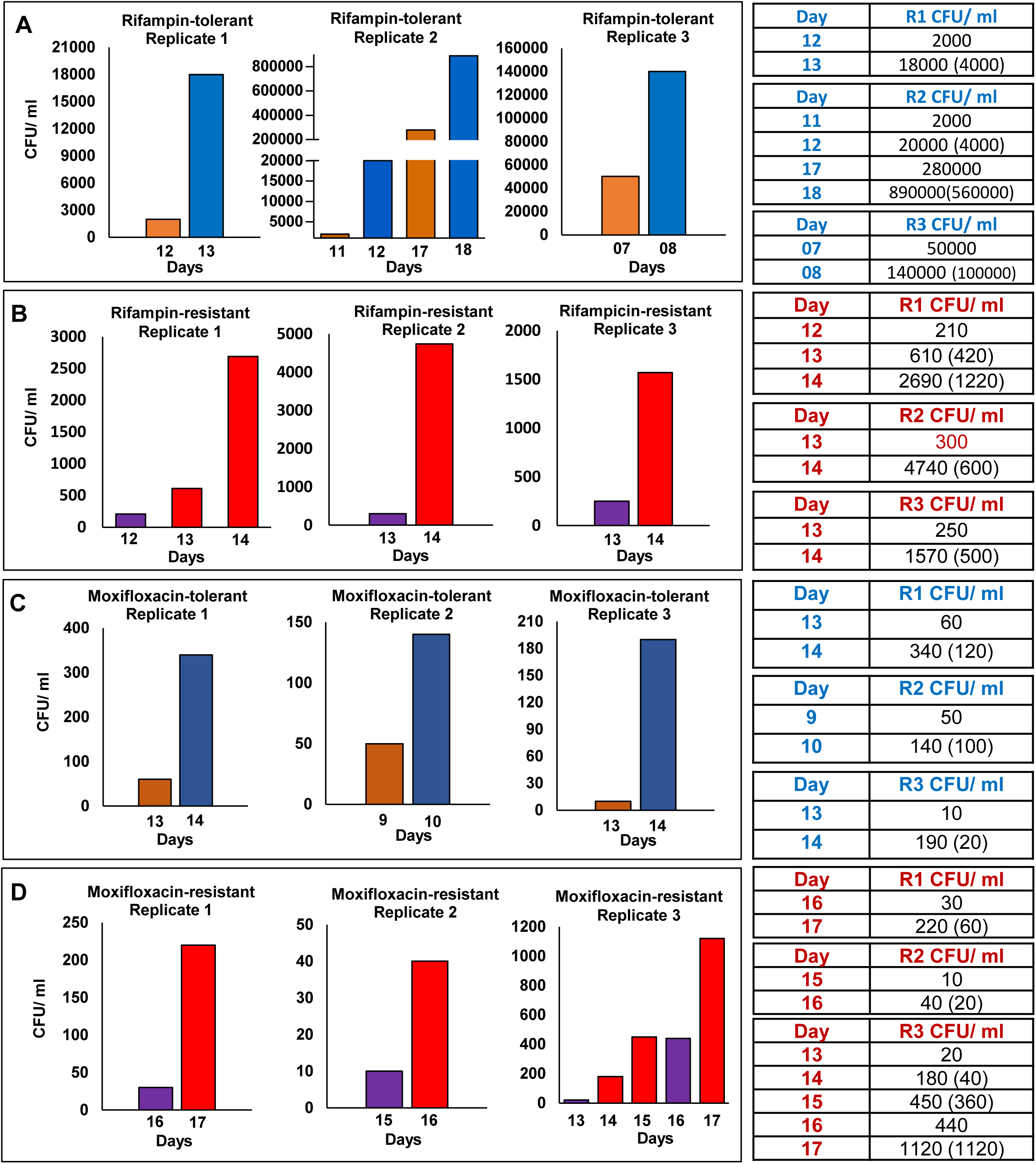
Response of *Mycobacterium tuberculosis* upon prolonged exposure independently to rifampicin and moxifloxacin. Observed CFU/ml (*expected CFU/ml given in parenthesis considering cell number doubling time as 24 hrs*) of *M. tuberculosis* cells, during antibiotic exposure to rifampin (1 µg/ ml; 10x MBC) or moxifloxacin (1 µg/ ml; 2x MBC), at every 24 hrs from antibiotic-free and antibiotic-containing [5 µg/ ml rifampicin (50x MBC) or 2 µg/ ml moxifloxacin (4x MBC)] Middlebrook 7H10 agar plates. CFU/ml of rifampin-tolerant cells, compared to the previous time point, was presented in **(A)** and of the rifampin-resisters was presented in **(B)**. Similarly, CFU/ml of moxifloxacin-tolerant cells, compared to that of the previous time point, was presented in **(C)** and that of the moxifloxacin-resisters was presented in **(D)**. The cfu values corresponded to the Fig. 1A (graph for rifampin-exposed *M. tuberculosis* cells) and Fig. S2 (graph for moxifloxacin-exposed *M. tuberculosis* cells) from our earlier work (Sebastian et al., 2017), where the unusual spurt in the cfu went unnoticed and unreported by us. Only the time points showing cfu spurt, taken from our earlier published work, have been re-presented here (Sebastian et al, 2017; Copyright @ American Society for Microbiology, Antimicrobial Agents and Chemotherapy, 61, 2017, e01343-16, https://doi.org/10.1128/AAC.01343-16).

### The *Msm* cells regrowing from the persister population contain multiple nucleoids

The unexpected unusually high fold-increase in the cfu during the regrowth phase could be due to the formation of multiple septae and multiple nucleoids in mother cells followed by their multiple division to generate multiple sister-daughter cells. Another possibility could be mono-nucleoid mother cells undergoing division with lesser mass and cell number doubling time (faster growth and division). A combination of both these phenomena also could cause the high spurt in cfu. We presumed that the second possibility of the cells undergoing faster growth and division is less likely to happen, as sudden change in the generation time has not been reported till date in bacterial systems.

Staining of MLP cells with the DNA-specific dye, Hoechst 33342, showed that 99.60 ± 0.61% of the cells at 0 hr (i.e. before antibiotic exposure) were having ≤2n nucleoid content and only 0.39 ± 0.61% cells showed >2n nucleoid content (n = 84) (**Fig. 5 A-D & Q**). On the contrary, 83.48 ± 3.91% cells from the regrowth phase (96 hr) showed >2n nucleoid content, with only 16.51 ± 3.91% cells having ≤2n nucleoid content (n = 896 cells) (**Fig. 5 E-P & R**). More images of the multi-nucleoid cells from the 96 hr of exposure are given in **Fig. S2**. It indicated that the *Msm* cells regrowing from the rifampin persister population were carrying multiple nucleoids, clearly indicating multiple nucleoid divisions, giving scope for future multiple divisions through multiple septation between pairs of nucleoids to generate multiple sister-daughter cells.

**Figure 5.**
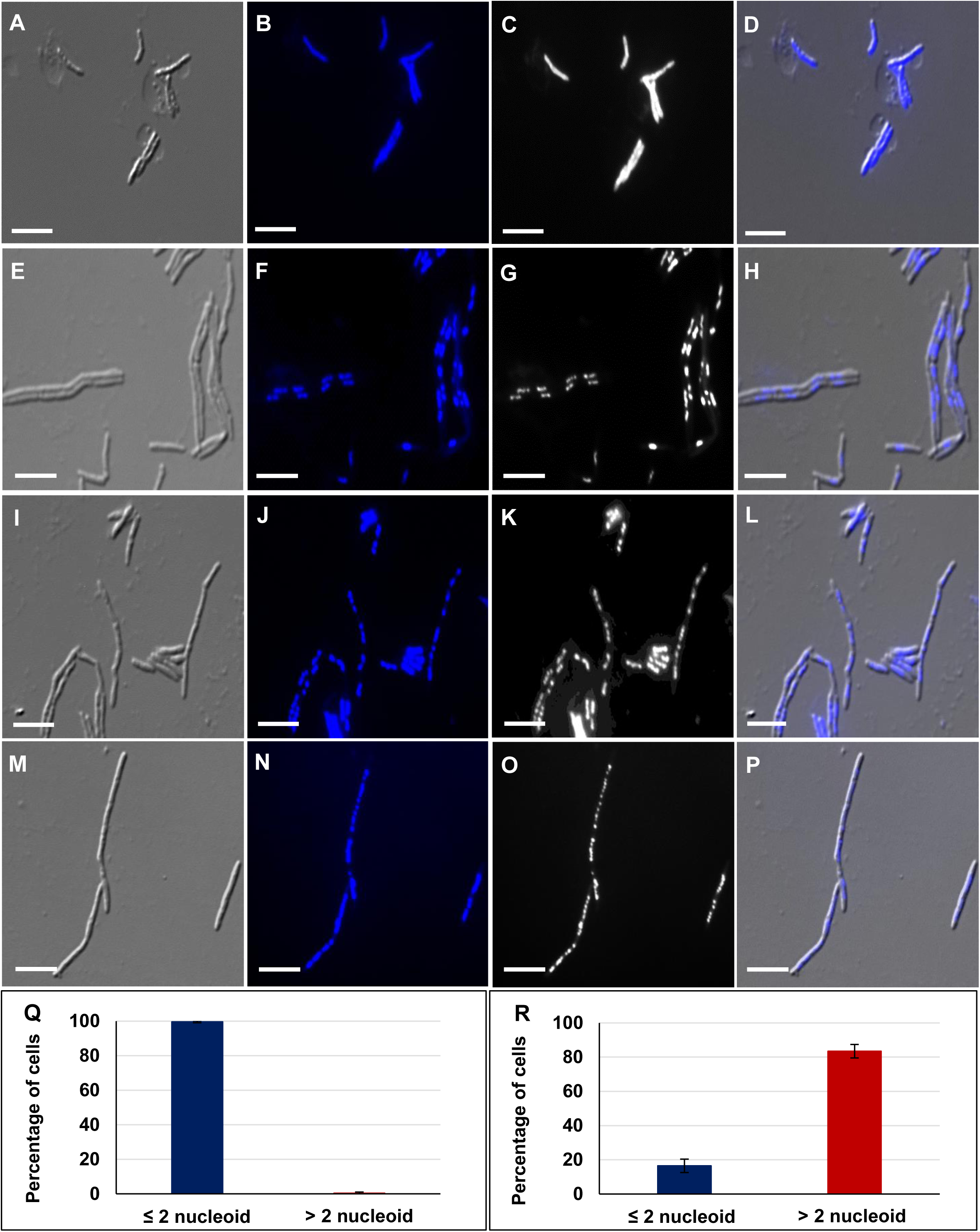
Hoechst staining of *Msm* MLP cells and the cells regrowing from rifampin persister population. **(A-D)** MLP cells prior to rifampin exposure (0 hr). **(E-P)** Cells (96 hr) regrowing from the persister population during antibiotic exposure. DIC images of: **(A)** MLP cells and **(E, I, M)** regrowing cells. The Hoechst fluorescence of the nucleoid profile of: **(B)** MLP cells and **(F, J, N**) regrowing cells. The bright field profile of the nucleoid profile of: **(C)** MLP cells and **(G, K, O)** regrowing cells. The DIC-Hoechst fluorescence merged images of **(D)** MLP cells and **(H, L, P)** regrowing cells. **(Q, R)** Quantitation of the cells: **(Q)** prior to rifampin exposure **(R)** 96 hr of rifampin exposure. n = 896 cells. Scale bar 5 µm.

### The *Msm* cells regrowing from the persister population possess multiple septae besides multiple nucleoids

Transmission electron micrographs of the cells, pertaining to 60 hr, 66 hr, 72 hr, 78 hr, 84 hr, 90 hr and 96 hr, regrowing from the persister population showed mostly elongated cells with multiple condensed nucleoids or multiple septae or with both positioned along the length of the cells and, at times, even at the poles (**Fig. 6A-H**). We could also observe the presence of shorter-sized cells with mid/polar septum (**Fig. 6I, J**). Among the total number of cells at 96 hr, 13.47 ± 3.41% of cells were septated cells and the remaining 86.52 ± 3.41% did not possess septum (n = 1081) (**Fig. 6Q**). Further, among the total number of cells, 23.39 ± 9.88% possessed multiple septae as compared to 76.03 ± 9.88% cells with single septum (n = 1081) (**Fig. 6A-C and D, I, respectively; 6R**). The atomic force micrographs of 96 hr cells regrowing from the rifampin persister population also revealed the presence of multiple ridges, each of which might be indicative of a septum beneath, which may be sites for multiple divisions in future (**Fig. S3**), as predicted (Dahl, 2004; Eskandarian *et al*., 2017).

**Figure 6.**
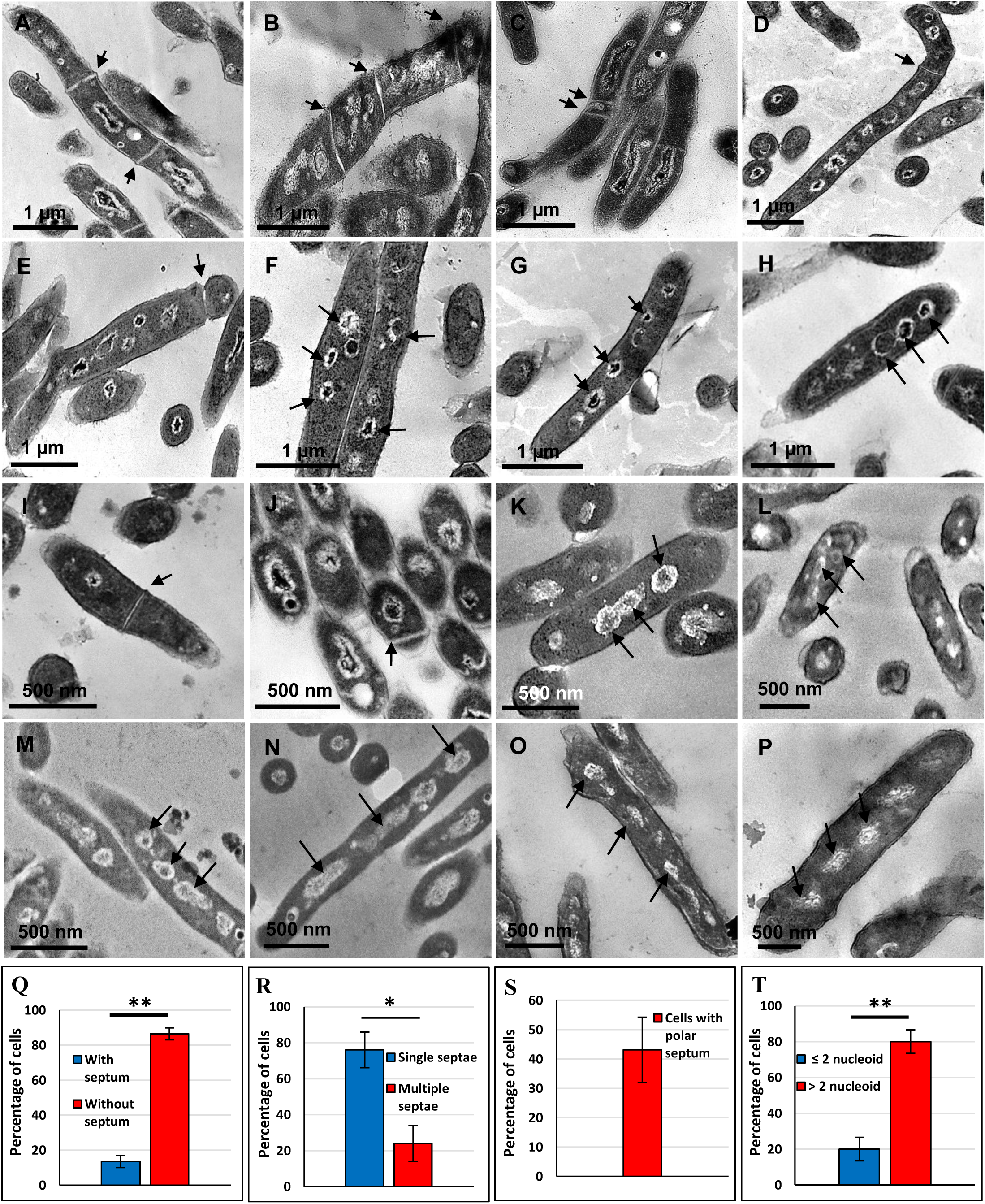
Transmission electron micrographs and quantitation of the *Msm* cells, regrowing from persister population, taken at different time points during rifampin exposure. **(A-H)** Elongated cells taken at 96 hr; **(I, J)** Shorter-sized (∼ 1 µm in length) cells at 96 hr; **(K-P)** Regrowing cells taken at: **(K)** 60 hr, **(L)** 66 hr, **(M)** 72 hr, **(N)** 78 hr, **(O)** 84 hr, and **(P)** 90 hr. **(Q-T)** Quantitation of the cells with / without septum, with nucleoid / septae. Most of the cells contain multiple nucleoids and multiple septae. High level of heterogeneity in terms of multiple nucleoids, multiple septae, position of the septae, and size and morphology of the cells could be noted. Arrow heads indicate nucleoid/septae. n = 1081 cells for **Q, R, T** and n = 210 septated cells for **S**.

Among the septated cells (n = 210 septated cells), 43.09 ± 11.14% of the cells were having polar septum owing to which seemingly anucleated cells also could be observed in the transmission electron micrographs (**Fig. 6E, J, S**). Further, among the total number of cells, 79.95 ± 6.55% cells were having >2n nucleoid content while the remaining 20.04 ± 6.55% possessed ≤2n nucleoid content (n = 1081) (**Fig. 6A-H, T**). More images of the cells with multiple nucleoids at 96 hr are given in **Fig. S4A-H**.

Shorter-sized cells (∼1 µm in length) with mid/polar septum could also be observed (**Fig. S4 I-K**). Genomic DNA profile obtained from the cells from six different colonies taken from the 96 hr sample, showed intact genome, thereby confirming lack of genome fragmentation in the cells (**Fig S4 L**). Thus, the intact genomic DNA isolated from the 96 hr cells suggested that the cells with multiple nucleoid-type images should not be misconstrued as cells with fragmented nucleoids. Further, nucleoid fragmentation was ruled out as it would not have given the unusual unexpectedly high spurt in the cfu during division as the cells with fragmented nucleoid would be unviable as single cells and hence would not generate progeny cells.

### Cellular heterogeneity in the regrowing population

Among the cells in the regrowing population, a high level of heterogeneity in terms of cells with different number of nucleoids, different number of septae, differences in the septal position, differences in lengths and morphology could be noted. We examined whether such level of heterogeneity was developed over a period of the growth and division of the cells in the regrowing population. The ultrastructural images of the cells in the regrowing population, at every 6 hrs during the regrowing phase from 60 hr to 90 hr, showed progressive increase in cell-length with increase in nucleoid content (**Fig. 6K-P**). Some of the cells showed elongated nucleoids (**Fig. 6M, N**), like in the bacterial cells that undergo nucleoid replication and segregation (Lin *et al*., 1997; Webb, 1997; Van helvoort *et al*., 1998; Ben-Yehuda *et al*., 2003; Errington *et al*., 2005; Zhang *et al*., 2009; Fisher *et al*., 2013; Gorle *et al*., 2017). In bacterial karyokinesis during cell division, the cells elongate to twice their length to form the sister-daughter cells and accommodate the replicating and segregating nucleoids (Sargent, 1975; Donachie *et al*., 1976; Donachie and Begg, 1989; Sharpe *et al*., 1998). Further, repeated karyokinesis (nucleoid replication and segregation) without cytokinesis (septation) causes elongation or filamentation of the cells (Adler *et al*., 1968; Spratt, 1975; Moya *et al*., 1998, Weiss *et al*., 1999; Rosenberger and Finlay, 2002; Chauhan *et al*., 2006; Justice *et al*., 2006; reviewed in Justice *et al*., 2008; Moller *et al*., 2013; Muckl *et al*., 2018). Therefore, with ∼3-4 µm as the average length of a non-dividing *Msm* MLP cell (Vijay *et al*., 2014b), formation of multiple nucleoids necessitated increase in cell-length anywhere from ∼8 µm (∼twice their average cell-length) to 20 µm (∼5 times their average cell-length), as observed from the images. From the cell-length of ∼3-4 µm of the cells at 0 hr of exposure (MLP cells), the cell-length progressively increased till 84 hr and reached a maximum by 96 hr (**Fig. 7; Fig. S5**). The phenotype observed in the DIC images correlated with the progressively elongated phenotype found in the transmission electron micrographs of the cells from the earlier hours of exposure to 96 hr (compare **Fig. 7, Fig. S5** with **Figure 6K-P**). By 102 hr, a sudden drop in the cell-length, to almost equal to that of the cells at 0 hr (MLP cells), was observed (**Fig. 7A, B and Fig. S5**). This drop might be indicative of the division of the multinucleated, multi-septated cells into mononucleated individual sister-daughter cells.

**Figure 7.**
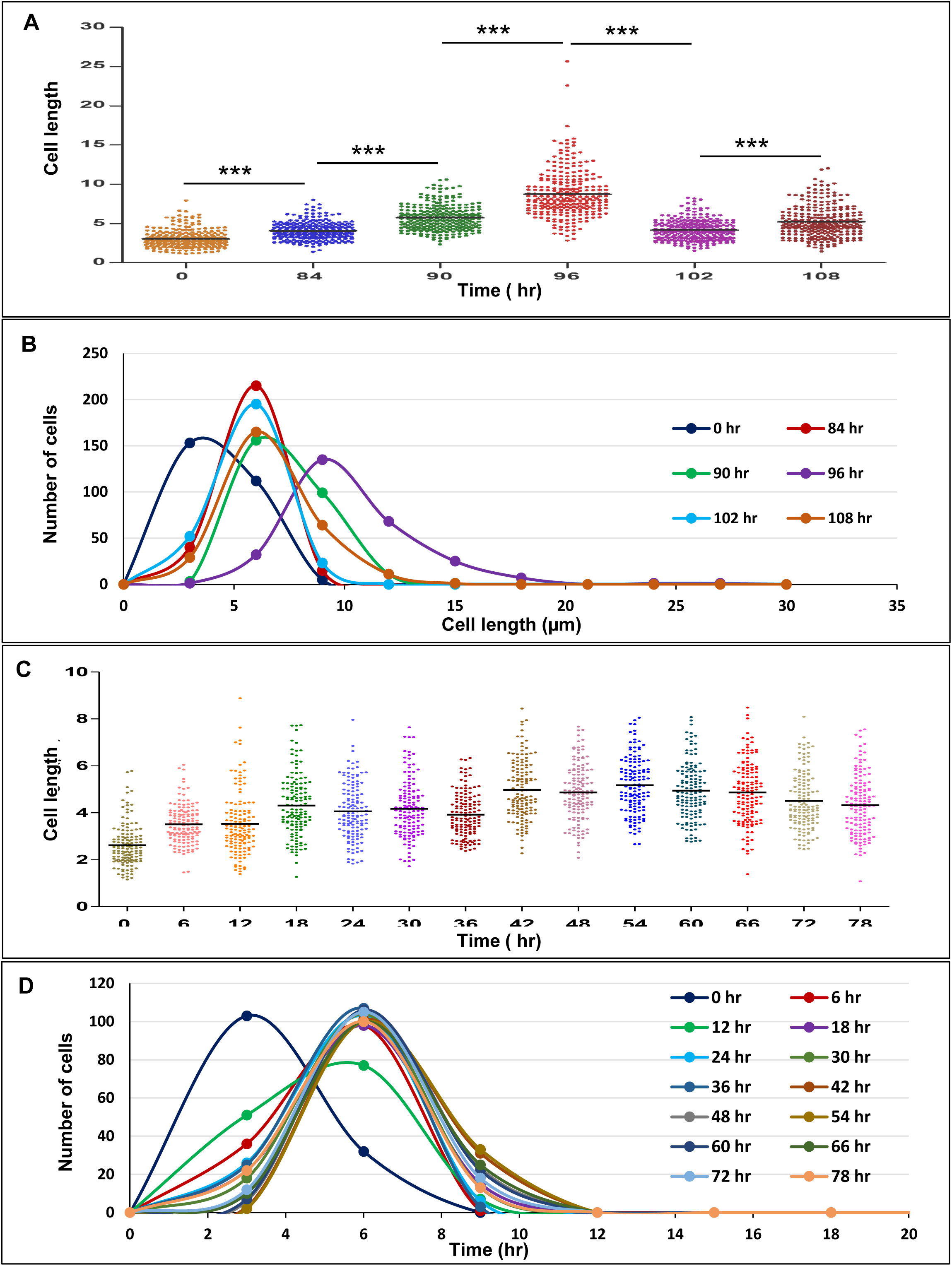
The determination of cell-lengths and cell-length distribution of the *Msm* cells during the entire period of exposure to rifampin. **(A)** The cell-lengths and their distribution among the cells: **(A, B)** in the late regrowth phase (84 hr to 108 hr), in comparison to those of the cells at 0 hr (MLP cells); **(C, D)** during the entire period of 0 hr to 78 hr, spanning the killing (0 hr to 36 hr), persistence phase (36 hr to 54 hr) and early regrowth phases (54 hr to 78 hr) of exposure to rifampin (n = 1890).

### No nutritional stress during the late hrs of regrowth

During the late hrs of growth, wherein stationary phase related nutritional stress sets in, the *Msm / M. bovis* BCG cells have been found to be shorter in size (Smeulders *et al*., 1999; Thanky *et al*., 2007; Markova *et al*., 2012; Wu *et al*., 2016; Dow and Prisic, 2018). Therefore, the drop in the cell-length of the cells in the 102 hr culture might be due to nutritional stress induced change into shorter size. However, the levels of glycerol in the medium up to 102 hr were comparable to its levels in the beginning of the culture, with very little levels utilised, indicating that there was no depletion of glycerol levels, which might have otherwise caused nutritional stress and consequential reduced-size phenotype (**Fig. S6**). The low levels of glycerol utilisation was expected as the entire duration of the culture from the start (0 hr) to 54 hr involved killing phase followed by the persistence phase where there would not have been any growth or division of the cells in the continued presence of rifampin (see **Figure 1A**). Subsequently, during the late regrowth phase from 102 hr, the glycerol levels steeply decreased, which correlated with the increase in the OD600 nm indicating glycerol utilisation for cell growth and division (**Fig. S6**). These observations ruled out the possibility of any nutritional stress which otherwise would have been suspected to be the cause for the reduction in the size of elongated cells into normal-sized cell size at 102 hr of exposure. Thus, the reduction in the length of the cells at 102 hr, as compared to the elongated cells at 96 hr (see **Fig. 7A, B, Fig. S4**), seemed most likely to be due to the division of the multi-nucleated, multi-septated cells into mononucleated individual sister-daughter cells.

### Time-lapse live-cell imaging of multiple constriction

Multiple septation and division was further monitored in live cells using time-lapse imaging of the multiple constriction of the cells from 90 hr till 102 hr of antibiotic exposure in order to confirm the observed phenomenon of the formation of multi-septated, multi-nucleated cells. Multiple constriction could be clearly observed among the regrowth phase cells wherein the daughter cells, which were generated by multiple constriction, again underwent multiple constriction and produced more than two sister-daughter cells in single division (**Fig. 8 and Video S1**). Tracking the division lineage of the cells showed multiple constriction of mother cells and of the sister-daughter cells that were produced from the multiple-constricted mother cells (**Fig. S7 traced from Video S1**).

**Figure 8.**
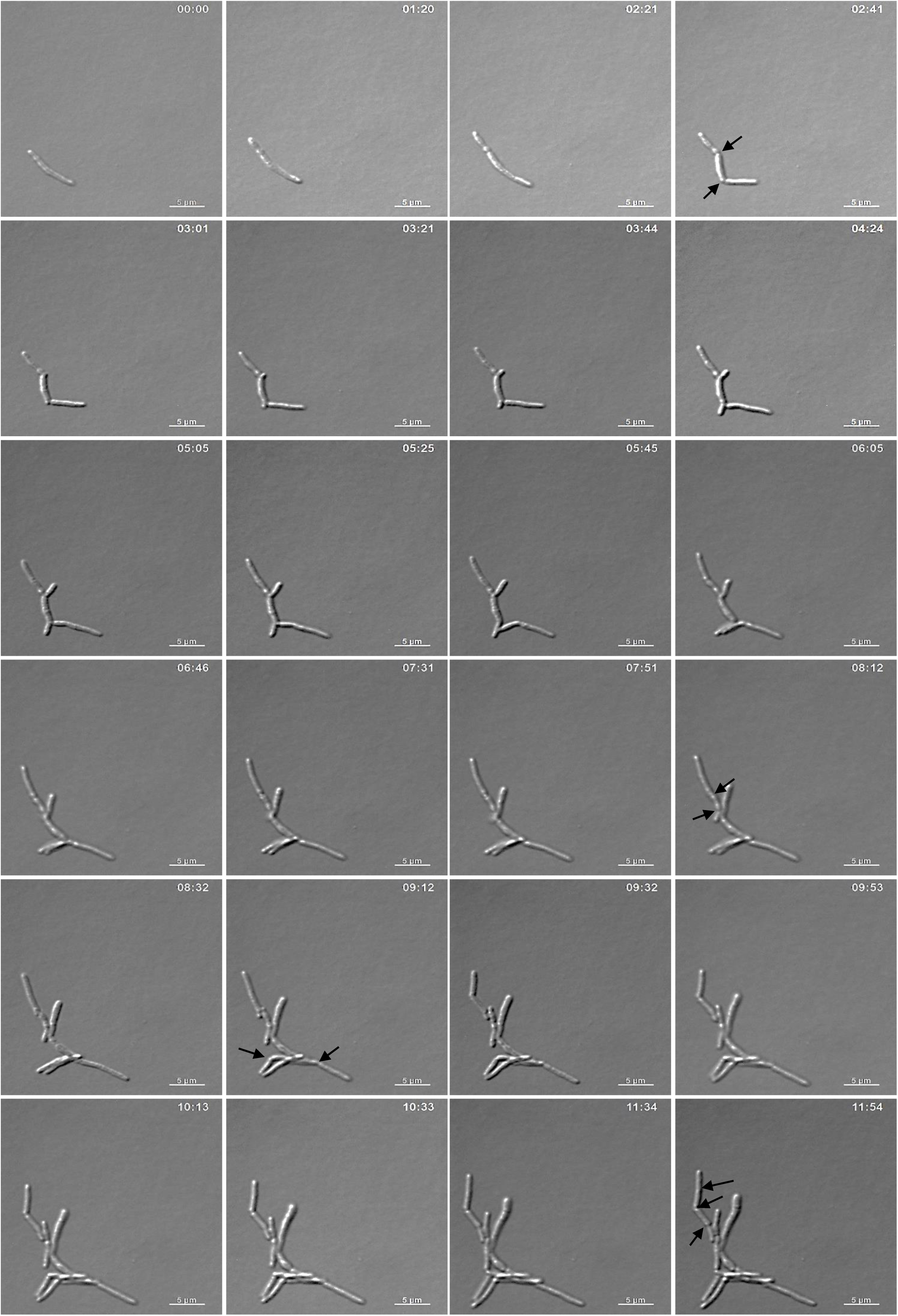
Live-cell time-lapse images of the cells from 96 hr regrowth phase. The cells show multiple constriction. Arrowhead indicated consriction. The images were from video S1.

### Regrowth phase cells were rifampin-resisters with mutation at RRDR

Regrowth and division in the presence of antibiotic normally occurs in the cells that have gained antibiotic resistance through ROS-mediated mutagenesis (Dwyer *et al*., 2009; Kohanski *et al*., 2010; Li *et al*., 2015; Sebastian *et al*., 2017; Hoeksema *et al*., 2018; Jin *et al*., 2018). Therefore, we wanted to find out whether the observed multiple division, resulting in sudden increase in the cfu of the regrowth phase population, was brought about by the rifampin-resister population carrying RRDR genetic mutation or by the rifampin-tolerant population lacking RRDR mutation. For this purpose, the colonies from the 84 hr, 90 hr, 96 hr and 102 hr of the regrowth phase from the rifampin-free plates were patched onto rifampin-containing plate (3x MBC; 125 µg/ml rifampin). All the colonies from the rifampin-free plates from the four time points grew on rifampin-containing plate, indicating that all the colonies from the regrowth phase had come from the rifampin-resistant mutants (**Fig. 9A**). This provided evidence that the cells with multiple septation might have been the resister population.

**Figure 9.**
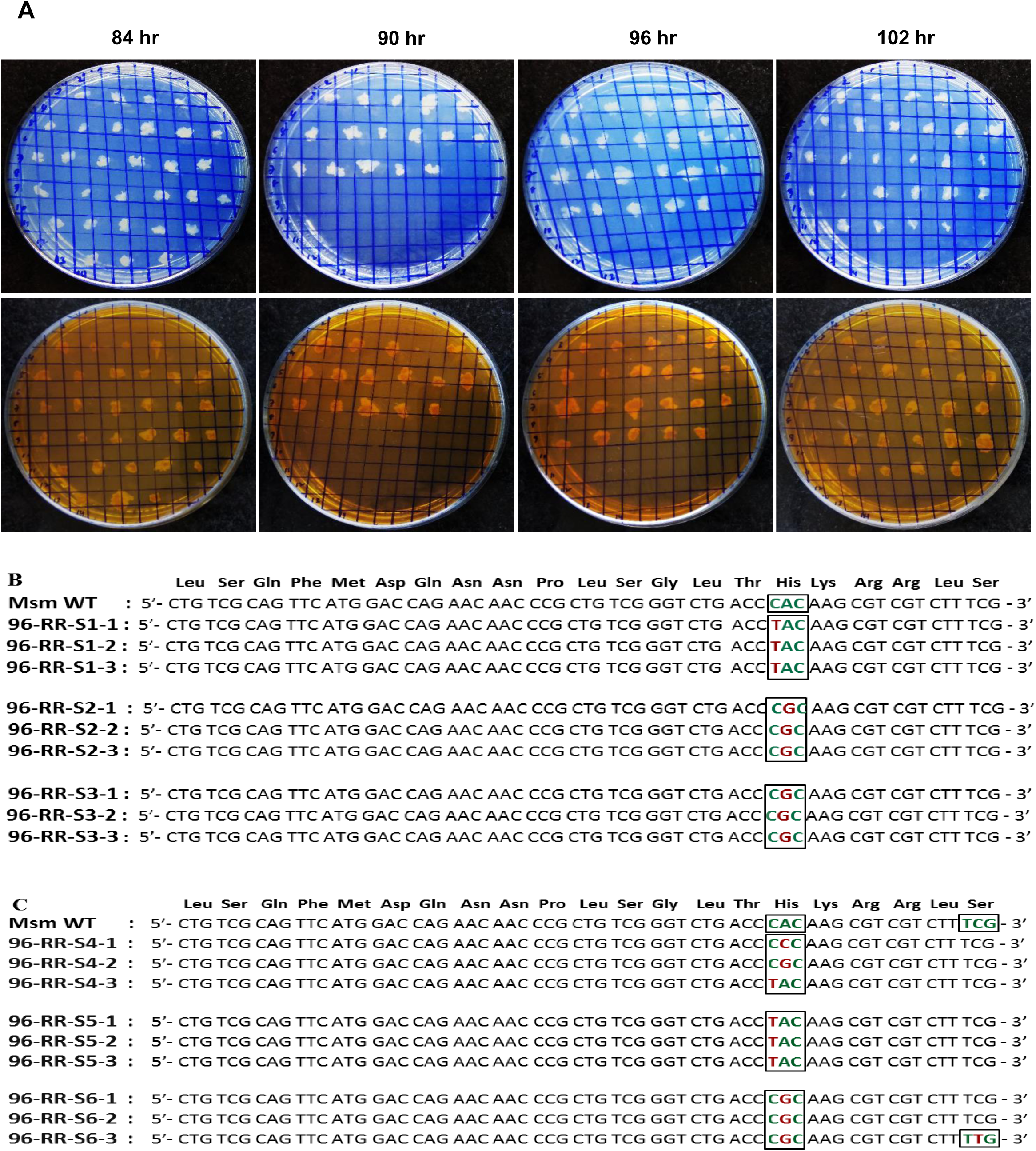
Colony patching of the cells taken from rifampin-free plate from different time points during the regrowth phase into rifampin-containing plate. **(A)**. Colonies, taken from the 84 hr, 90 hr, 96 hr and 102 hr from the rifampin-free master plates, were patched onto rifampin-free plates (top panel in blue colour) and rifampin (125 µg/ ml) plates (lower panel in orange colour). **(B, C)**. Comparison of the RRDR sequence of the wild-type genome with the RRDR sequence of the cells from the: **(B)** rifampin plates and **(C)** rifampin-free plates. In both **(B)** and **(C)**, n = 3 experimental samples, each with biological replicates.

Although the cells had grown on rifampin plates, there was a possibility that they could be phenotypic or genotypic resisters. Sequencing of the RRDR of the genomic DNA from the 96 hr colonies from the rifampin plates, performed using RRDR primers (**Table S1**), showed mutations in all the colonies, confirming that these were indeed the cells that were carrying genetic mutation at the RRDR of the *rpoB* gene, which conferred resistance to rifampin (**Fig. 9B**). Sequencing of the colonies, from the rifampin-free plate, which were originally obtained from rifampin-containing liquid culture, also possessed mutation in the RRDR (**Fig. 9C**). This confirmed that the observed mutations were already present in the cells in the rifampin-containing liquid culture itself and were not generated on the selection plate containing rifampin.

We and other groups have shown the generation of reactive oxygen species (ROS) in the antibiotic-exposed *Mtb* cells (Grant *et al*., 2012; Piccaro *et al*., 2014; Nandakumar *et al*., 2014) and in the rifampin/moxifloxacin-exposed *Mtb* persister cells (Sebastian *et al*., 2017) and *Msm* persister cells (Swaminath, 2017). Consistent with this findings, we could detect high levels of hydroxyl radical specific hydroxyphenyl fluorescein (HPF) fluorescence from the persister phase cells (48 hr onwards) (**Fig. S8**). The HPF fluorescence steadily decreased till 90 hr, during the emergence of rifampin-resistant mutant cells in the regrowth phase, and subsequently plateaued (**Fig. S8**). It indicated the generation of hydroxyl radical in the persister phase cells and the decline of the same in the regrowth phase cells once the cells had gained resistance, and become free from the antibiotic stress, as reported by us for rifampin-exposed *Mtb* cells (Sebastian *et al*., 2017) and moxifloxacin-stressed *Msm* cells (Swaminath, 2017). The reciprocal decline/rise nature of the profile of cfu and HPF fluorescence indicated that the HPF fluorescence, indicative of hydroxyl radical generation, was highest when the cells were in the persister phase, and that the hydroxyl radical generation declined when the cells started regrowth and division after gaining relief from the rifampin stress by acquiring rifampin resistance through RRDR mutation (**Fig. 9B, C and Fig. S8**).

### Naturally formed rifampin-resisters do not show sudden spurt in cell division

It may be recalled here that it was the rifampin-resistant mutants, which emerged from the persistence phase through the multiple division of multi-septated, multi-nucleated elongated cells, showed the phenomenon of sudden abnormally high spurt in division. Therefore, we wanted to verify whether the phenomenon would be shown by the rifampin-resistant mutants, which arise naturally from the mid-log population and not from the rifampin persister population. For this purpose, we checked whether the rifampin-resistant genetic mutants, which were selected from the mid-log population against rifampin by direct plating, and therefore not emerged from persister population, would show sudden increase in the cfu when re-exposed to rifampin for prolonged duration. All the rifampin-resisters, which were selected by plating the MLP culture on 3x MBC rifampin plate, possessed mutation in the RRDR (**Fig. S9A**). Upon culturing these mutants in 25 µg/ml rifampin-containing liquid medium and plating on antibiotic-free plate, the cells showed the expected 2-fold increase in the cfu within 3 hrs of cell number doubling time. It indicated that unlike the rifampin-resistant mutants formed from the persister population, the rifampin-resistant mutant cells from the MLP culture underwent normal cell division without any spurt (**Fig. S9B, C**). Thus, the abnormally high spurt in the cell division was a unique cell division behaviour of the ROS-induced rifampin-resistant mutants from the rifampin persister population only.

### Heterogeneity of regrowing cells prevents molecular analysis of the mechanism

In order to understand the molecular mechanism that drives the formation of multi-septated, multi-nucleated cells and consequential sudden spurt in cell division, we isolated and purified total RNA of high integrity from the regrowing population of cells from the 96 hr time point and performed quantitative RT-PCR. The genes selected for the quantitative PCR included those involved in cell division, DNA repair, error-prone DNA polymerisation, SOS regulon, oxidative stress response, and few other controls. However, we got widely varying results from multiple biological replicates of the total RNA (data not shown). We presume that this variation could be due to the cellular heterogeneity of the regrowing population revealed by the TEM images which showed wide differences in the proportions of the cells with multiple septae, multiple nucleoids, polar septum, dividing and non-dividing cells. Hence the total RNA isolated from this mixture of different types of cells is quite likely to contain specific mRNAs but in different proportions, giving wide variations in the quantitative RT-PCR data. We found the fractionation of these different subpopulations difficult.

## Discussion

### Bacteria metamorphose into multi-septated and/or multi-nucleated phenotypes under diverse stress conditions

In response to diverse stress conditions, bacteria of diverse genera, including mycobacteria, have been found to stop division, grow to become multi-septated and/or multi-nucleated, and regrow back with a spurt in cfu to establish mutant population. For example, carbon starvation of *M. smegmatis*, *Mycobacterium fortuitum*, and *Mycobacterium peregrinum* under saline conditions triggered the formation of small resting cell (SMRC) morphotype, which developed into multi-septated, multi-nucleated cells that divided to generate mononucleated SMRCs (Wu *et al*., 2016). Similarly, starved large resting cells (LARCs) also metamorphosed into multi-septated, multi-nucleated cells. In another study, although both the *M. tuberculosis* clinical strains, TB 282 and TB 284 of smooth and rough morphology, respectively, showed loss of viability upon recovery from exposure to ambient air drying, TB 284 showed a sudden spurt in cfu from 60 to 90 min during the recovery of the sample (Klein & Yang, 2014). It was not clear whether the TB 284 cells, which showed spurt in cfu, were multi-septated and/or multi-nucleated. However, one might infer that a sudden spurt in cell number could have most probably come from multi-septated and/or multi-nucleated cells.

Another example of external stress condition influencing ploidy levels is that of the cyanobacterial model strain, *Synechocystis* sp. PCC 6803. These cells show the ploidy levels of 53 copies and 35 copies at lower light intensity and at higher phosphate concentrations, respectively (Zerulla *et al*., 2016). Complete absence of phosphate induced a rapid reduction in the genome copy number, indicating that DNA replication ceased, and genomes were distributed to the daughter cells. However, prolonged incubation of the stationary phase cultures in the absence of phosphate led to the formation of monoploid cells. Similarly, the haloarchaeal species, *Halobacterium salinarum, Haloferax mediterranei*, and *Haloferax volcanii*, are all polyploid, with most of them having a higher copy number during the exponential growth phase than in the stationary phase (reviewed in Zerulla & Soppa, 2014). The advantages of polyploidy were suggested to be low mutation rate, high resistance to x-ray and desiccation. *H. volcanii* was found to use genomic DNA both as the genetic material and as a storage for phosphate in polymeric form. Thus, under phosphate starvation, *H. volcanii* significantly decreases ploidy, thereby enabling cell multiplication, but reducing the genetic advantages of polyploidy. However, it is not known whether the polyploidy cells were multi-septated.

### Phenotypic plasticity as a strategy for survival under stress conditions

Increasing number of studies have shown that morphological plasticity involving formation of multi-nucleated, multi-septated filamentation by bacterial cells under stress conditions might be a common strategy for survival (reviewed in Justice *et al*., 2008). Stress conditions such as high pressure (Zobell & Cobet, 1964), nutrient depletion (Pine & Boone, 1967), DNA damage from SOS response (Radman, 1975; Bos *et al*., 2012), host innate immune reactions (Justice *et al*., 2006), and reduced water content (Stackhouse *et al*., 2012) have also been found to induce multi-nucleated and/or multi-septated filamentation on bacterial cells. An earlier study on 1.1 OD600 nm *Mtb* cells using scanning electron microscopy (SEM) showed the presence of cells with multiple ridges (Dahl, 2004). It was suggested that the cells with multiple ridges might be having multiple septae just below the ridges, indicative of the presence of multi-compartments carrying multi-nucleoid cells. It is possible that the cells might have been approaching stationary phase as the phenomenon was observed in a 1.1 OD600 nm *Mtb* cells. A decade later, atomic force microscopy (AFM) of *Msm* cells undergoing division demonstrated that the septum position would have the morphological mark of ridge-with-trough-on-both-sides type of undulating surface corresponding to the future site of cell division (Eskandarian *et al*., 2017).

### Sudden spurt in cfu of mycobacterial systems

The response of triggering an increase in cfu against antibiotics has been noted in mycobacteria as well. For instance, upon treatment of *M. tuberculosis* H37Rv infected nude mice with rifampin and INH, the cfu per lung showed an initial sudden fall from ∼10^8^ to ∼10^5^ at 2 months post-treatment. But, surprisingly, the bacillary load sharply increased to 10^6^ cfu per lung on the 3^rd^ month and to 10^8^ cfu per lung on the 4^th^ month, with a subsequent plateau (Zhang *et al*., 2011). It was of interest to note that the bacilli recovered from the increased cfu were INH-resistant. However, it was not mentioned whether the cells that underwent spurt in the cfu were multi-septated or multi-nucleated. As could be seen from our data, since all the stressed and mutated cells in the population would not undergo spurt in division, it is difficult to categorically state whether the 10^8^ cfu per lung on the 4^th^ month (which arose from 10^5^ cfu per lung on the 2^nd^ month) included cells that divided in spurt. Nevertheless, the response and the behaviour thereof of *M. tuberculosis* against antibiotics in mice lung and in *in vitro* culture are comparable in terms of the sudden increase in the cfu. This in turn validates the present *in vitro* study indicating that the response of the bacilli with a sudden spurt in cell division is an inherent trait of the bacilli irrespective of their growth habitat and pathogenic/non-pathogenic status.

### SOS response leading to mutagenesis as a strategy to regrow under stress conditions

Bacteria generate SOS response against many stress agents, including antibiotics, leading to the generation of genome-wide mutations confer mutations which in turn becomes a strategy against succumbing to lethality inflicted by the stress agents (Ryan, 1959; Radman, 1975; Witkin, 1976; Hude *et al*., 1990; Kato *et al*., 1994; McKenzie *et al*., 2000; McKenzie *et al*., 2001; Miller *et al*., 2004; Asad *et al*., 2004; Janion, 2008; Peng *et al*., 2011; Rodríguez-Beltran *et al*., 2012; Kreuzer, 2013; Baharoglu *et al*., 2014; Qin *et al*., 2015; Handel *et al*., 2015; Dapa *et al*., 2017; Sebastian *et al*., 2017; Rodríguez-Rosado *et al*., 2018; Chistyakov *et al*., 2018; Crane *et al*., 2018; Hoeksema *et al*., 2018; Geisinger *et al*., 2019; Swaminath, 2017). When a multi-septated, multi-nucleated filamentous bacterial cell acquires SOS-driven mutations, which can confer resistance to the stress agent, it can initiate synchronous division along the entire length of the filament, generating large numbers of viable, normal-sized, sister-daughter cells, as reported (Stackhouse *et al*., 2012). Therefore, if the SOS-driven mutation happens to be in the antibiotic target gene, then that mutant would get selected when confronted with the antibiotic.

In a study on the response of *E. coli* to subminimal inhibitory concentrations of ciprofloxacin, the exposure converted rod-shaped bacteria into multi-nucleated filamentous phenotype (Bos *et al*., 2015). Upon continuous maintenance of ciprofloxacin at low levels, the filamented cells began to divide asymmetrically and repeatedly from the filament tips, generating budded ciprofloxacin-resistant progeny cells. The mutant chromosomes were produced due to the mutagenic SOS response and probable recombination of the new alleles between chromosomes, which in turn might have conferred improved survival advantage. The successful segregation of multiple mutant chromosomes toward the filaments’ tips was found to favour the birth of antibiotic-resistant cells. The authors proposed that the multi-nucleated filamentation was a precursor to the generation of highly resistant bacteria since it provided a strategy for the generation of mutant chromosomes that may confer survival advantages, as proposed (Bos *et al*., 2015). In the light of the conclusions made in this study on the antibiotic-exposed *E. coli*, the multi-septated and multi-nucleated phenotype and acquiring of mutations by the persistence phase population of the *Msm* cells exposed to rifampin for prolonged duration in our study indicate that the morphological and genetic changes might be to ensure the survival of the rifampin-exposed cells.

### Multi-nucleation/septation, mutagenesis and spurt in cell division in antibiotic-stressed mycobacteria

In the background of all these observations, our present study on the rifampin/moxifloxacin-exposed *Msm* persister and regrowth cells and the earlier study on the rifampin/moxifloxacin-exposed *Mtb* persister and regrowth cells (Sebastian *et al*., 2017) showcase the phenomenon of sudden spurt in cell division shown by the antibiotic-resistant mutant cells emerging from the antibiotic persister population in the context of the phenomenon of antibiotic persistence. The data from fluorescence microscopy, AFM, TEM and live-cell imaging of the cell samples from 90 hr to 96 hr confirmed that regrowth phase cells possessed multiple septae with multiple nucleoids. These multi-nucleated, multi-septated elongated mother cells underwent multiple constrictions to give rise to multiple sister-daughter cells. This, in turn, could explain the observed sudden increase in the cfu of the cells from the regrowth phase to unexpected levels. Live-cell imaging, which showed multiple constrictions of multi-nucleated, multi-septated cells into mononucleated individual sister-daughter cells having lengths comparable to those of individual nondividing MLP cells, supported that the reduction in cell-length, from the elongated phenotype of the 96 hr cells to shorter size cells of the 102 hr, was due to multiple divisions of the parent cells.

Since large number of cells regrowing from the rifampin persister cells were found to be multi-nucleated, it is important to rule out the possibility of the multi-nucleated cells being cells with fragmented nucleoids. Had he multiple nucleoids been fragmented pieces of nucleoids, then such fragmented nucleoids: (i) would not have shown nucleoids with clear boundaries, (ii) would be of different shapes and sizes unlike a perfect same-sized round shape, and (iii) would have been found dispersed or diffused throughout the cell but would not be in an arranged manner. Apart from the absence of all these characteristics of fragmented nucleoids, the very clear sharp bands of the genomic DNA obtained from the regrowing cells of 96 hr (see Fig. S4) showed that the genomic DNA of the cells regrowing from the rifampin persister population were not degraded.

In addition to the multi-nucleated cells, negligible proportion of the cells in the regrowing population showed anucleated portion due to polar septation. Anucleated cells had been found in ParD mutant strains of *E. coli* (Hussain *et al*., 1987) and *spo0J* mutants of *Bacillus subtilis* (Ireton *et al*., 1994). Overproduction of ParA in mycobacteria also generated elongated cells with multiple nucleoids without septum, and cells with asymmetric septum with nucleoids in both sister-daughter cells, indicating defects in cell division (Maloney *et al*., 2009). But the overproduction of ParB resulted in the formation of anucleated cells with asymmetric septum, with nucleoid in one of the sister-daughter cells only (Maloney *et al*., 2009). It was possible that specific gene systems might be getting activated or repressed when the persister cells gained mutations and began to regrow. However, the persister cells generated elevated levels of hydroxyl radical, which is a sequence-non-specific mutagen (Sakai *et al*., 2006). Therefore, it was possible that the persister cells might have suffered genome-wide mutations, as we had shown in the rifampicin-exposed *Mtb* persister cells (Sebastian *et al*., 2017). In such a scenario, there would be every possibility for the mutations occurring on several genes, including Par genes, involved in chromosome partitioning. Such mutations in turn might have been the reason behind the formation of negligibly low proportions of anucleated cells with polar septum, multi-nucleated cells with only very few septae and so on, generating heterogeneity.

### Are there differences between the cells regrowing from the rifampicin/moxifloxacin persisters and isoniazid persisters?

In the background of our study on the physiological mechanisms that drive the generation of rifampin/moxifloxacin-resistant mutants from the respective persister population, it is important to compare it with an interesting study on the behaviour of *M. smegmatis* persister cells against isoniazid. The *M. smegmatis* persister cells against isoniazid were found to be in a dynamic state of balance between the number of cells that undergo division and cell death such that the net cfu would not change in the presence of isoniazid (Wakamoto *et al*., 2013). However, it would not be possible for non-tolerant or non-resister cells to proliferate in the presence of an antibiotic. Therefore, it could be inferred from the study on the response of *Msm* persister cells to isoniazid that most probably the dynamically growing and dividing *Msm* persister cells in the presence of isoniazid would have already gained either phenotypic or genotypic tolerance/resistance to isoniazid. Thus, compared to the behaviour of the *Msm* isoniazid persister cells in the presence of isoniazid, the behaviour of the rifampin and moxifloxacin persister cells of *Mtb* and *Msm* in the present study were very different.

### Broad implications of the present study

The fact that this behaviour was found in both the pathogenic *Mtb* and the saprophytic *Msm* cells upon prolonged exposure to two antibiotics of totally diverse mode of action, rifampin and moxifloxacin, confirmed the phenomenon to be a natural one that is neither specific to antibiotics nor to any specific mycobacterial species. This phenomenon of the formation of multi-septated polyploidy mycobacterial cells seems to be a unique mode of generation of the genetically resistant mutants against antibiotics, when they are freshly emerging from the antibiotic persister population. Further, since even *Escherichia coli* also showed similar response against subminimal inhibitory concentrations of ciprofloxacin (Bos *et al*., 2015), these findings have a broad significance as a general strategy adopted by diverse bacterial genera to emerge as drug-resistant strains against antibiotics.

Thus, many bacteria, including mycobacteria, use the strategy of sudden spurt in cell division, after gaining resistance against diverse stress conditions, either by generating multi-septated and/or polyploidy cells. However, whether the mycobacterial persister cells, formed in response to exposure to microbicidal concentrations of antibiotics, would respond in a similar manner through the formation of multi-septated and multi-nucleated cells undergoing spurt in the generation of antibiotic-resistant genetically mutant progeny cells remained unknown till the present study. Since we had observed the phenomenon in *Mtb* cells also, possession of such a trait by the tubercle bacilli in the patients can pose a major impediment in TB treatment for the eradication of drug resistance developed by the pathogen.

## Materials and methods

### Bacterial strain and culture conditions

*Mycobacterium smegmatis* mc^2^155 (Snapper *et al*., 1990; *Msm*) was grown in autoclaved sterilised fresh Middlebrook 7H9 liquid cultures containing 0.2% glycerol (Fisher Scientific) and 0.05% Tween 80 (Sigma) under continuous gyration at 170 rpm, 37°C by taking 5:1 head space to volume ratio. Plating experiments were performed on Mycobacteria 7H11 agar plates (Difco) with and/or without rifampin (125 µg/ml; MP Biomedicals) at 37°C for 3 to 4 days. *Msm* cultures were also grown in the presence of moxifloxacin (Cayman Chemicals) at 0.5 µg/ml (3.759 X MBC), prepared from the stock of 2 mg/ml solution in DMSO. All the experiments were conducted using actively growing mid-log phase cells with an approximate cell density of 10^6^ cells/ml.

### Growth curve experiments

From the starting of the experiment, liquid cultures were exposed to 25 µg/ml concentration of the antibiotic rifampin (MP Biomedicals). The antibiotic stock solution (50 mg/ml) was made by dissolving rifampin powder (MP Biomedicals) in DMSO (Merck Millipore) and filter sterilised using 0.22 µm PVDF syringe filters (Millex-GV). Dilutions of the antibiotic (rifampin) was performed using 0.22 µm PVDF (Millex-GV) filter sterilised DMSO (Merck Millipore). Similarly, moxifloxacin (0.5 µg/ml; 3.759x MBC) was added to an independent 100 ml *Msm* secondary culture. Post-antibiotic exposure, cells were taken at every 6 hr time interval and plated on antibiotic free and antibiotic containing (125 µg/ml) Mycobacteria 7H11 agar (Difco) plates to check the actual number of colony forming unit (cfu) present at that particular time point and their respective resisters. Dilution of the cultures for the plating was performed using fresh Middlebrook 7H9 media. Plates were incubated in Innova 4200 Incubator Shaker at 37°C for 3-5 days.

### Fold increase in the cfu calculations

Fold increase in cfu at any specific time point was calculated by normalising the cfu value of that specific time point with the cfu obtained at its previous time point.

### Hoechst staining of nucleoid

200 µl of the cells from the mid-log as well as from regrowth phases were taken into an Eppendorf tube and pelleted down at ∼ 5000 × g for 10 min at room temperature. The supernatant was removed, and the cell pellet was washed once with fresh Middlebrook 7H9 (Difco) media and pelleted again with same centrifugation conditions. The cell pellet was resuspended in 200 µl of fresh Middlebrook 7H9 (Difco) media and 10 ng/ml concentration of the Hoechst 33342 stain (Sigma) was added. This mixture was incubated at 37°C (Innova 4200 Incubator Shaker) for 10 min in dark. After 10 min, the cells were washed to remove the excess dye by pelleting down at ∼ 5000 × g for 10 min at room temperature and the pellet was resuspended in fresh Middlebrook 7H9 (Difco) media and a drop of the cell suspension was taken on a clean glass slide and dispersed evenly. The slide was air dried at room temperature for 10 min in dark and a small drop of glycerol was added and covered with coverslip. The slide thus made was observed under Carl Zeiss AXIO Imager MI microscope at 100x by putting a drop of immersion oil on top of the coverslip.

### Sample preparation for Atomic Force Microscopy

Sample preparation for atomic force microscopy was carried out as described (Bolshakova *et al*., 2001) with minor modifications. In brief, 1 ml of the cells from regrowth phase (96 hr) were taken in a fresh Eppendorf tube and harvested by centrifuging them at ∼ 5000 × g for 10 min at room temperature. The cell pellet was washed once with fresh Middlebrook 7H9 liquid media and parallelly a glass coverslip was cleaned once with acetone (Merck Millipore), and once it dried, a drop of the cell suspension was added on the coverslip. The cover slip was air dried for 10 min at room temperature and post air dry, it was washed gently with fresh de-ionised water and again set for drying for 10 min at room temperature. The cells fixed on the glass coverslip were observed under Atomic Force Microscope in contact independent mode of scanning.

### Sample preparation for Transmission Electron Microscopy

Cells from regrowth phase (96 hr) were taken and processed for transmission electron microscopic analysis as described (Takade *et al*., 1983; Vijay *et al*., 2012) with minor modifications. In brief, 1 ml of the cells from regrowth phase (96 hr) were taken into Eppendorf tube and harvested by centrifuging them at ∼ 5000 × g for 10 min at room temperature. Supernatant was discarded and the cell pellet was resuspended in 0.15 M sodium cacodylate-HCl buffer pH 7.4, containing 1% osmium tetroxide (OsO4) (w/v) (Sigma) and incubated (Innova 4200 Incubator Shaker) for 1 hr at room temperature in dark. After 1 hr, cells were washed with the same buffer by pelleting them at ∼ 5000 × g for 10 min at room temperature. The cell pellet was resuspended in 0.15 M sodium cacodylate (w/v) (Sigma)-HCl buffer pH 7.4 containing 2% glutaraldehyde (v/v) (Sigma), 2% tannic acid (w/v) and incubated at room temperature for 2 hrs in dark. The cell suspension was washed with the same buffer for 10 min at room temperature at ∼ 5000 × g and the pellet was resuspended in 0.15 M sodium cacodylate buffer containing 1% OsO4 and incubated overnight at 4°C. Post overnight incubation, the cell suspension was washed with the same buffer at ∼ 5000 × g for 10 min at room temperature and the pellet was subjected for dehydration with serial dilutions of ethanol (Merck Millipore) (20%, 30%, 50%, 70%, 90%, 100%, and 100%). After dehydration, the cell pellet was resuspended in 50% LR White resin (1:1 volume of LR White resin (London Resin Company) and 95% ethanol (Merck Millipore)) and the cell suspension was stored in 4°C till further use. Gelatine blocks were prepared by spinning down this cell suspension at room temperature for 10 min at ∼ 5000 × g and an aliquot of the cell pellet was taken and made it to adhere at the bottom of the gelatine capsule (Electron Microscopy Sciences) and then capsule was filled with 100% LR White resin. This gelatine capsule was kept at 60°C for 48-56 hrs till it became rock hard. Finally, the blocks were trimmed and cut into 70 nm ultra-thin sections using ultra microtome (Power Tome XI). These sections were collected on a copper grid (150 mesh × 165 µm pitch; Sigma-Aldrich) and the copper grids containing sections were subjected for uranyl acetate and led citrate staining. The stained sections were observed at 120 kV under Transmission Electron Microscopy (Tecnai Bio-TWIN).

### Cell length measurements and size distribution graph

DIC images of cell before (0 hr) and post antibiotic exposure (till 120 hr) were captured using Carl Zeiss AXIO Imager M1 microscope and measured using AxioVision 4 software.

### Glycerol estimation

Free glycerol concentration in the culture supernatant was measured by converting glycerol to formaldehyde using periodate (White *et al*., 1974) and spectrophotometric estimation of formaldehyde by means of Hantzsch reaction at 410 nm (Nash 1953; Bondioli and Bella, 2005). Glycerol concentration in the supernatant was measured at different time points, as described (Bondioli and Bella, 2005), with minor modifications. In brief, the reagents were prepared as follows. (i). acetic acid stock solution: 1.6 M aqueous solution in autoclaved distilled water; (ii). ammonium acetate stock solution: 4.0 M aqueous solution in autoclaved distilled water; (iii). 0.2 M acetyl acetone solution: 200 µl of acetyl acetone was added into 5 ml of acetic acid stock solution, mixed well, and 5 ml of ammonium acetate stock solution was added; (iv). 10 mM sodium (meta) periodate solution: 21 mg of sodium (meta) periodate was dissolved in 5 ml of acetic acid stock solution, and once sodium (meta) periodate was completely dissolved, 5 ml of ammonium acetate stock solution was added. Working solvent: equal volumes of autoclaved distilled water and 95% ethanol.

Different concentrations of glycerol standards were prepared as 0.4, 0.2, 0.1, 0.05, 0.025, 0.0125, and 0.00625% by serial dilution using distilled water. 13 µl of glycerol standard was dissolved in 117 µl of distilled water. 390 µl of working solvent was added to 130 µl of the diluted standard. Three hundred and twelve (312) µl of sodium (meta) periodate solution was added and mixed on a vortex mixer for 30 sec. Subsequently, 312 µl of 0.2 M acetyl acetone solution was added and kept in a heating block at 70°C for 1 min. The sample was then immediately cooled by keeping the tube in ice. Absorbance was measured immediately at 410 nm in a UV-Visible spectrophotometer. Standard calibration curve was constructed and equation for the straight line, i.e. y = mx + c, was used to determine the glycerol concentration in the culture medium, where y is absorbance at 410 nm, m is the slope, c is the y intercept, and x is the unknown concentration of glycerol to be determined as x = (y-c) ÷ m.

The glycerol concentration in the culture supernatants of the antibiotic exposed culture was determined. In brief, culture supernatants at different time points were collected by centrifuging 1 ml of *Msm* culture at ∼5,000 × g for 10 min at room temperature, from independently grown biological triplicate cultures. Samples were treated with sodium (meta) periodate and acetyl acetone solution in the same way as in the case of the standard. Absorbance was measured at 410 nm and the glycerol concentration (x) was calculated with the equation of the straight line of the standard calibration curve.

### Patch plating of cells from rifampin-free plate to rifampin-containing plate

A small portion of the colony from antibiotic free plate was picked up with the help of toothpick and seeded on antibiotic (125 µg/ml rifampin; MP Biomedicals) containing plate. This plate was incubated at 37°C (Innova 4200 Incubator Shaker) bacterial incubator for 3-5 days.

### Expected and observed cfu

Fold change of cfu at any specific time point was calculated by considering the next time point cfu value as ‘B’ and subtracted this value with earlier time point cfu value (considered as ‘A’) and the resulting value was divided with the time point ‘A’. Expected cfu increase within 6 hrs will be 4 times the number of cells present at the previous time point. Thus, any time point where the cells showed more than 4-fold increase in cfu, was considered an unusual division. The table was made accordingly with the observed and expected cfu values. The time points where the cell number was higher than the expected cfu were highlighted in box.

### Live cell time-lapse microscopy

Live cell time-lapse microscopy of *Msm* regrowth phase cells was performed using agarose pad method, as described (Jong *et al*., 2011; Joyce *et al*., 2011; Vijay *et al*., 2014a, b) with minor modifications. Agarose pad was prepared by using spent media of 90 hr rifampin (25 µg/ml) exposed culture and 1.75% low melting point agarose on a clean glass slide. After solidification, a portion of the agarose (about 1/5^th^ of the total agarose pad area) was cut out using a blade to make a well for the introduction of rifampin into the agarose pad. Towards one side of the well, a tip was attached which was connected to a syringe for removal of rifampin containing medium. 10 µl of 90 hr rifampin (25 µg/ml) exposed *Msm* cells were placed on top of the agarose pad and spread evenly by tilting the slide. The slide was covered by a cover slip at 45° angle from the base glass slide, leaving a portion of the well open for the introduction of rifampin with a syringe. The agarose pad with the cells was kept at 37°C for 1 hr incubation to facilitate cell adhesion. This slide was observed under Carl Zeiss AxioVision 4 microscope using live cell imaging option, at 100x magnification (DIC), with 0.2 µm slice distance Z-stacking at 37°C. Individual cells were observed and the DIC images were taken at every 10 min time interval. The data was analysed and the parameters like cell length and cell constriction were determined on the images, using AxioVision 4 and ImageJ software.

### Cell pellet preparation for genomic DNA

Colonies taken from antibiotic free or antibiotic containing plates were resuspended in sterilised fresh Middlebrook 7H9 (Difco) liquid media and subjected for pipetting at least 10-15 times to remove the clumps. The cell suspension was re-inoculated into fresh sterilised Middlebrook 7H9 (Difco) liquid medium with and/or without antibiotic (25 µg/ml rifampin; MP Biomedicals) and incubated at 37°C (Innova 4200 Incubator Shaker) with a gyration speed of 170 rpm till the culture reaches an approximate cell density of 10^6^ cells/ml. Cultures (50 ml) were taken into Sorvall tube and harvested at ∼ 5400 × g at room temperature. Supernatant was discarded and the cell pellet was subjected for genomic DNA isolation.

### Genomic DNA isolation

Genomic DNA was prepared from mid-log phase as well as from rifampin-exposed cell cultures by phenol-chloroform extraction method. In brief, cell pellet was lysed by resuspending in 1 ml of Tris-HCl-EDA buffer (10 mM Tris-HCl containing 1 mM EDTA, pH 8) containing 3 mg/ml lysozyme (Chicken egg white; Fluka) and 1 mg/ml lipase (from *Candida cylindracea*; Sigma) in an Eppendorf tube. This cell suspension was kept at 37°C water bath for 4 hrs with intermittent mixing by inverting the Eppendorf tubes. The cell lysis was carried out by adding 2% SDS (Sigma) (final concentration) and incubated at 55°C dry bath for 10 min again with intermittent mixing (once in 30 min) by inverting the Eppendorf tubes. The cell lysate was centrifuged at ∼ 7800 × g for 10 min at 4°C. The supernatant (∼ 800 µl) was collected using cut tips and subjected for phenol-chloroform extraction using equal volumes of ice-chilled phenol (Tris-HCl saturated pH 7.8; Sisco Research Laboratories) chloroform (Nice Chemicals) mixture in 1:1 ratio (volume to volume). The suspension was mixed by inverting the Eppendorf tube ∼ 5 to 10 times and centrifuged at ∼ 7800 × g for 10 min at 4°C. The supernatant was collected and mixed with equal volume of phenol (Tris-HCl saturated; pH 7.8) and again mixed by inverting the Eppendorf tube ∼ 5 to 10 times. This suspension was centrifuged at ∼ 7800 × g for 10 min at 4°C. To this supernatant, equal volumes of ice chilled chloroform (Nice Chemicals) was added and mixed again by inverting the tubes for 5 to 10 times. This mixture was again subjected for centrifugation at ∼ 5000 × g for 10 min at 4°C and the supernatant was collected. To this supernatant, 1/10^th^ volume of 0.3 M sodium acetate (pH 7.4; Sigma) (final concentration) was added and mixed well with pipette. After this, volumes of 95% ice-cold ethanol (Merck Millipore) was added and mixed vigorously for 15 to 20 times by inverting the Eppendorf tubes. The suspension was kept at 4°C overnight for better precipitation. After overnight incubation at 4°C, DNA was pelleted down by centrifuging the suspension at ∼ 11300 × g for 20 min at 4°C. Supernatant was discarded and the cell pellet was washed once with 70% ethanol (Merck Millipore) and the pellet was air dried for 10 min. After 10 min, the obtained DNA pellet was dissolved in 1x Tris-HCl-EDTA buffer (10 mM Tris-HCl, 1mM EDTA, pH 8). This DNA was subjected for RNase A treatment by incubating with 1 µl of 10 mg/ml RNase A (bovine pancreatic RNase; Sigma) at 50°C for 60 min. After RNase treatment, DNA was re-extracted by phenol-chloroform method and precipitated as mentioned earlier. Finally, the obtained gDNA was dissolved in 1x Tris-HCl-EDTA buffer (10 mM Tris-HCl, 1mM EDTA, pH 8) and stored at 4°C till further use.

### PCR amplification of RRDR region

RRDR locus was PCR amplified using RRDR specific forward, and reverse primers (**Table. 4**) using Phusion Polymerase (Thermo Scientific, USA). The PCR amplified products were run on 1.5% agarose gel and the specific amplified band was eluted using gel elution kit (GeneJET Gel Extraction Kit; Thermo Scientific, USA). Sequence determination was performed for both the strands of the DNA by Chromous Biotech, Bangalore, India.

### HPF staining of *M. smegmatis* cells

HPF staining of cells post antibiotic exposure (25 µg/ml rifampin; Sigma) was carried out by taking 500 µl of the culture from respective time points into Eppendorf tubes and incubated them with 5 µM 3’-(p-hydroxyphenyl fluorescein (HPF; Invitrogen) (0.5 µl of HPF from 5 mM stock) (Setsukinai *et al*., 2003 and Mukherjee *et al*., 2009) at 37°C for 15 min with shaking at 170 rpm in a bacteriological incubator shaker (Innova 4200 Incubator Shaker) in dark. The cells, post incubation with HPF, were centrifuged at ∼ 5000 × g for 10 min at room temperature, resuspended in 200 µl of fresh Middlebrook 7H9 (Difco) and the cell suspension was taken into a clear bottom black multi-well Polystyrene assay plate. Fluorescence was observed using Tecan Infinite® 200 PRO series plate reader at the excitation maxima of 490 nm and emission maxima of 520 nm. The fluorescence obtained by multi-plate reader was divided with the cfu value obtained by plating to get the fluorescence value per cell. The obtained value was normalised with the value from the 48 hr time point and the fold increase in the fluorescence per cell was plotted in the graph against time.

## Acknowledgements

With highest respects and regards, PA dedicates this work as a tribute to Prof. T. Ramakrishnan (late), who led the pioneering, fundamental and foundation laying work on the biochemistry and molecular biology of *Mycobacterium tuberculosis* at Indian Institute of Science, Bangalore, India.

## Author’s contribution

PA, KJ designed the experiments; PA contributed reagents, materials, and analysis tools; KJ, DS, RRN, AvP, AP performed experiments; PA, KJ, DS, RRN, AvP, AP analysed data; PA, KJ wrote the manuscript; all authors have read the manuscript.

---

**Fig. S1.**
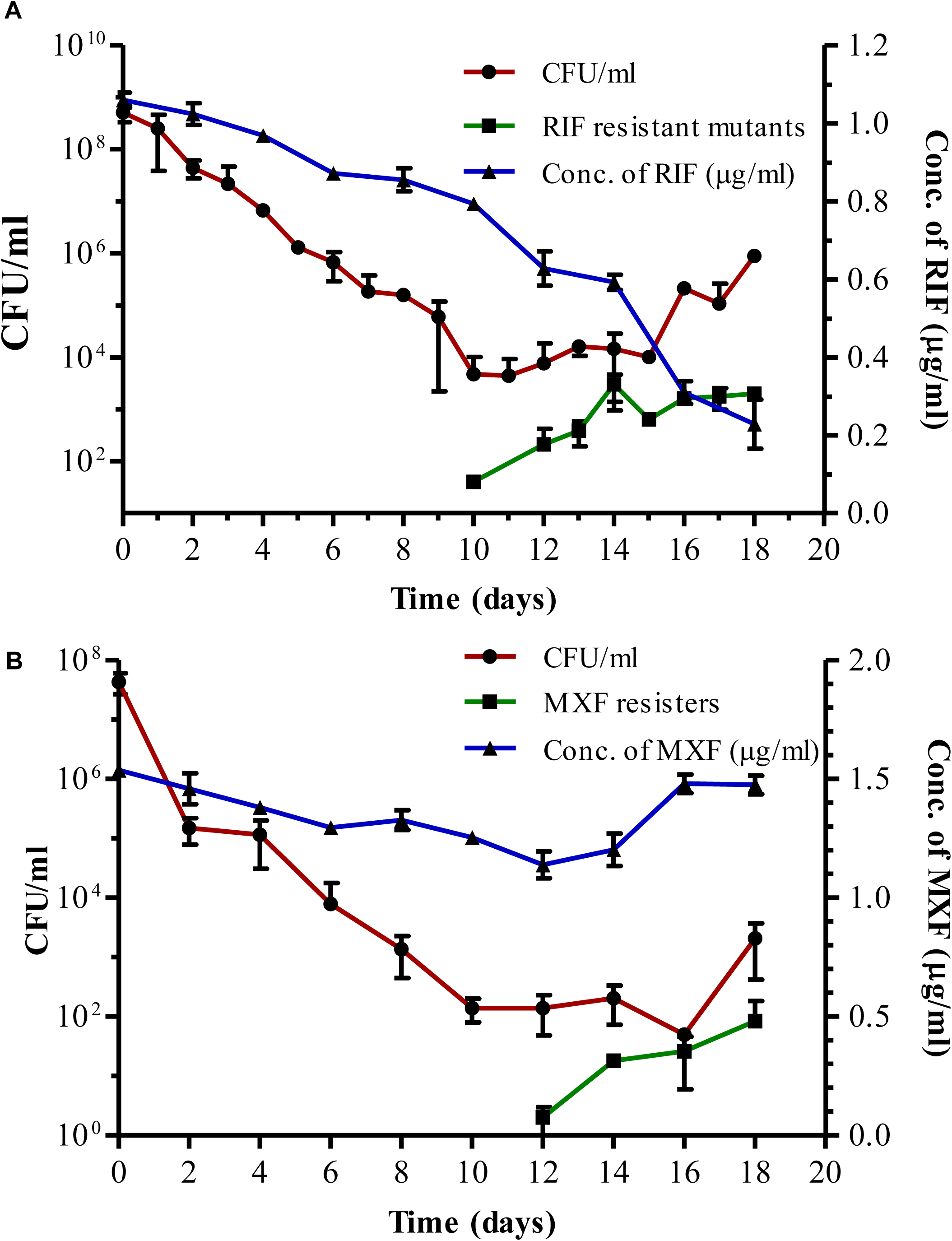
Susceptibility profile of *Mtb* cells to MBCs of rifampicin and moxifloxacin. **(A)** The susceptibility profile of *Mtb* cells, exposed to 10x MBC rifampicin (1 μg/ml) for 18 days, obtained by plating aliquotes of the culture on rifampicin-free plate every day (●; red line) and in parallel on rifampicin-containing (50x MBC) plates (▪; green line). The right y-axis shows the concentration of rifampicin during the exposure for 18 days (▴; blue line). **(B)** The susceptibility profile of *Mtb* cells, during exposure to 2x MBC moxifloxacin (1 μg/ml) for 18 days, obtained by plating aliquotes of the culture on moxifloxacin-free plate every day (●; red line) and in parallel on moxifloxacin-containing (4x MBC) plates (▪; green line). Right Y-axis shows the moxifloxacin concentration during the entire exposure period. Sebastian et al, 2017; Copyright @ American Society for Microbiology, Antimicrobial Agents and Chemotherapy, 61, 2017, e01343-16, https://doi.org/10.1128/AAC.01343-16.

**Figure S2.**
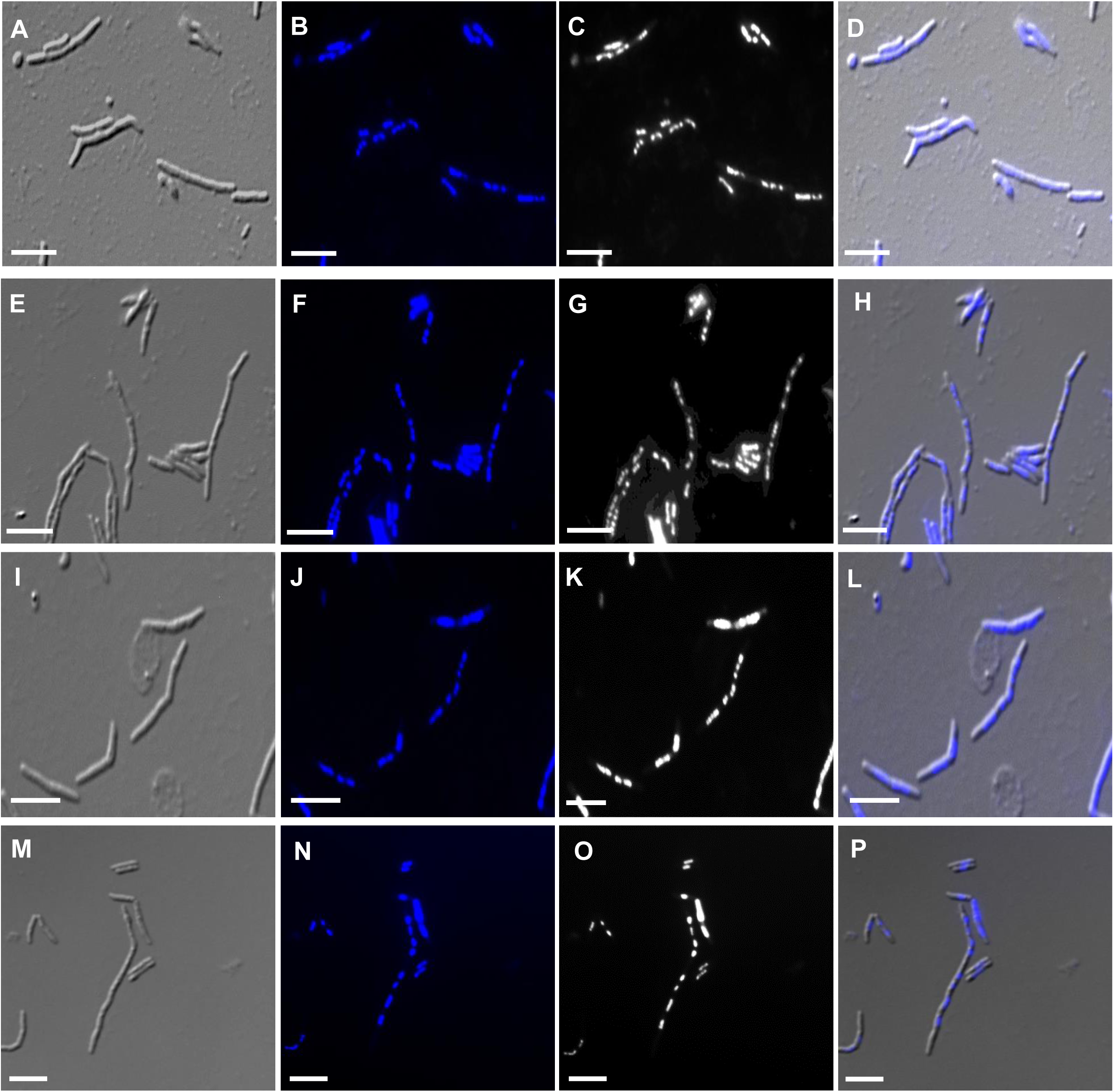
Hoechst staining of *Msm* cells regrowing from rifampin persister population. **(A-P)** More images of the cells (96 hr) regrowing from the persister population during antibiotic exposure. **(A, E, I, M)** DIC images; **(B, F, J, N)** nucleoid profile; **(C, G, K, O)** nucleoid profile in bright field; **(D, H, L, P)** merged images. Scale bar 5 µm.

**Figure S3.**
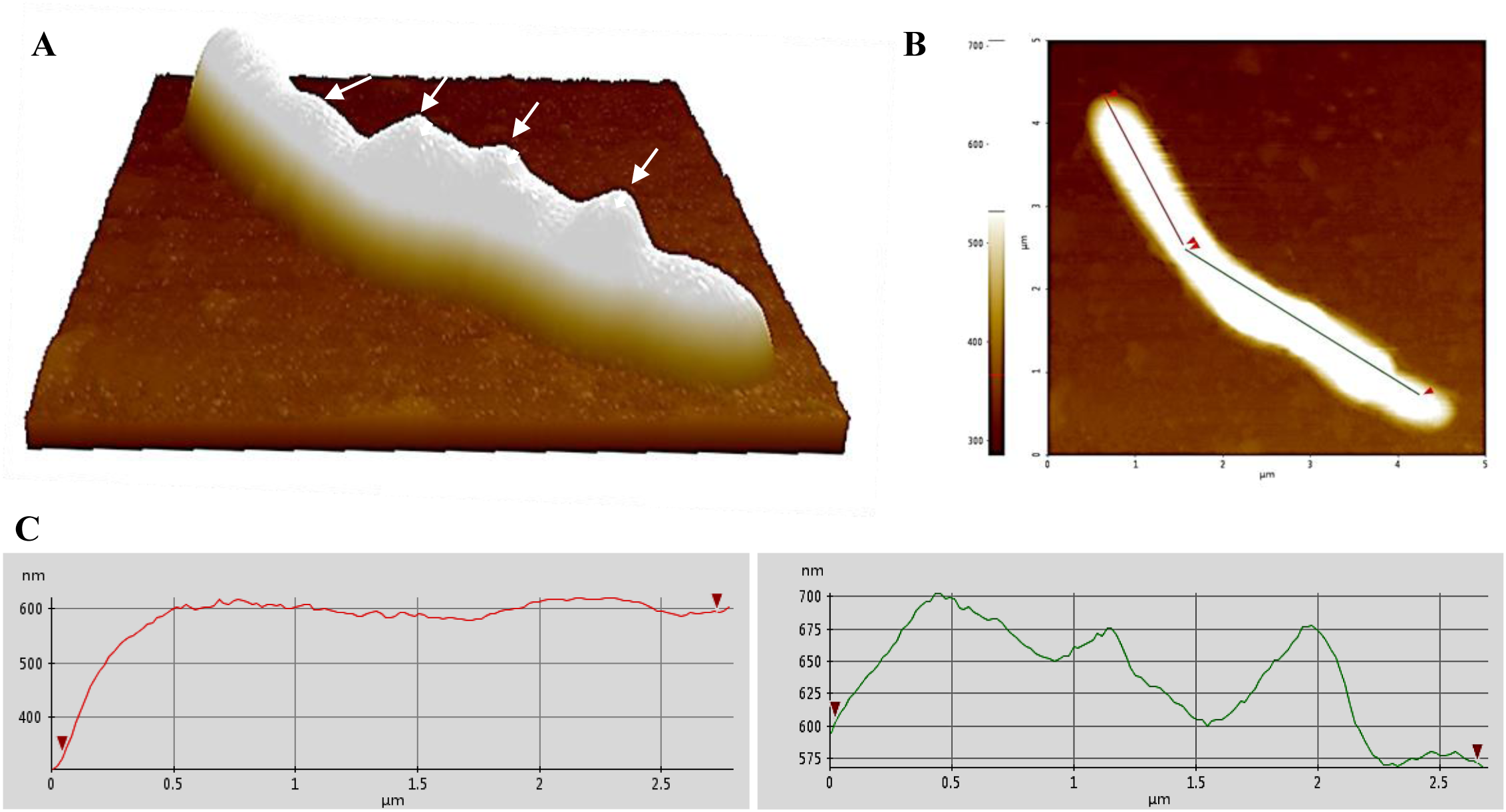
Atomic force micrographs of *Mycobacterium smegmatis* cells during rifampin exposure. **(A)**. 3D representative image of a cell taken from the 96 hr regrowth phase showing ridges-and-troughs type of cell surface. **(B)**. The flattened image indicates the area selected for making the line profiles (red and green). **(C)**. Line profiles (red and green) representing the ridges-and-troughs type of cell surface. Arrow heads indicate circular waveform troughs, probably corresponding to multiple septae beneath.

**Figure S4.**
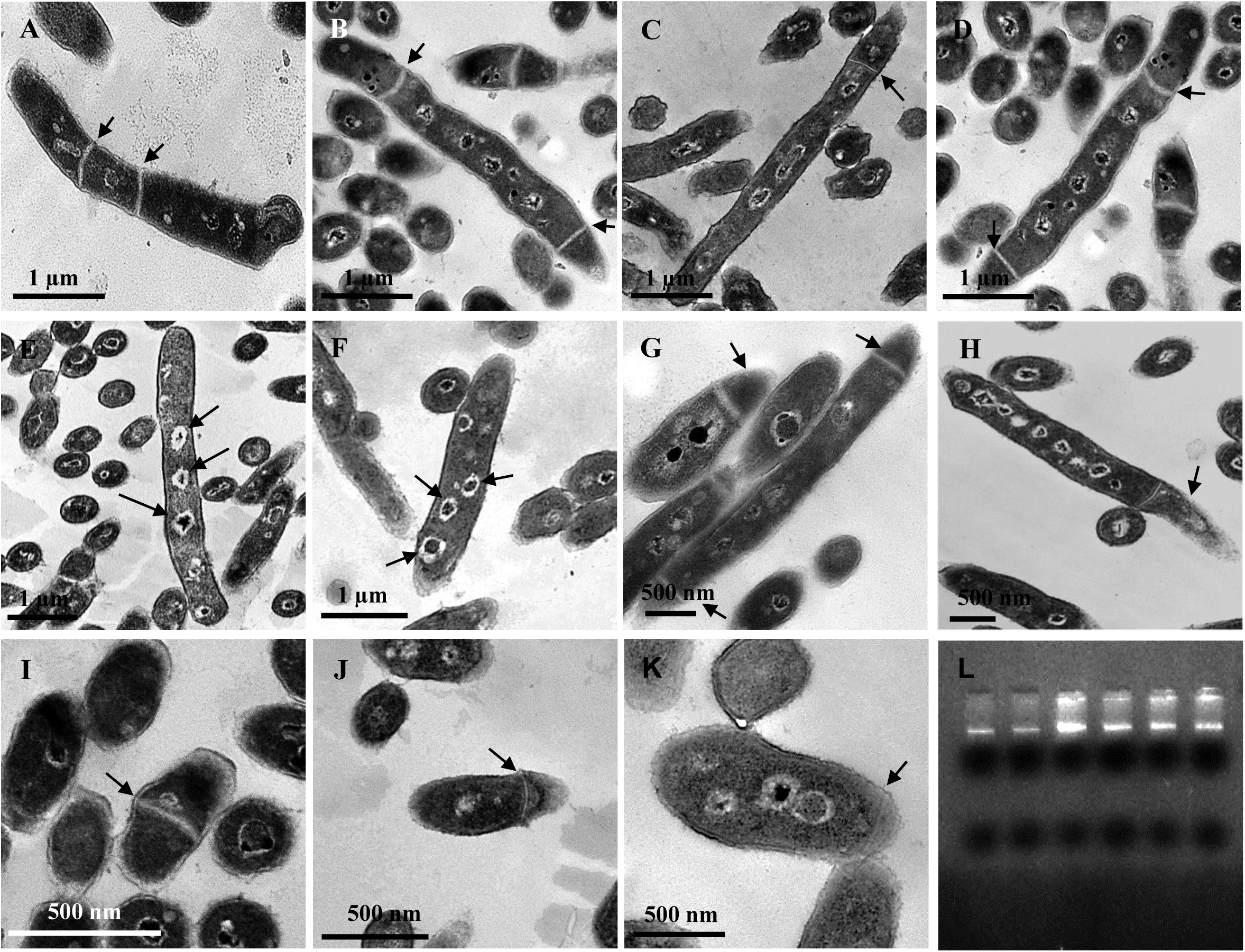
Transmission electron micrographs of the *Msm* cells, regrowing from persister population, at 96 hr. Cells with: **(A, B)** multiple septae, **(C, D)** polar septum with nucleoids, **(E, F)** multiple nucleoids. **(G, H)** Cells with anucleated portions due to polar septation. **(I-K)** Mid/polar septation in shorter-sized cells (∼1 µm in length). **(L)** Genomic DNA profile obtained from the cells from six different colonies taken from the 96 hr sample, showing intact genome, thereby confirming lack of genome fragmentation. A high level of heterogeneity in terms of multiple nucleoids, multiple septae, septal position, size and morphology could be noted. Arrow heads indicate nucleoid/ septae/ anucleated cell (n = > 1081 cells).

**Fig S5.**
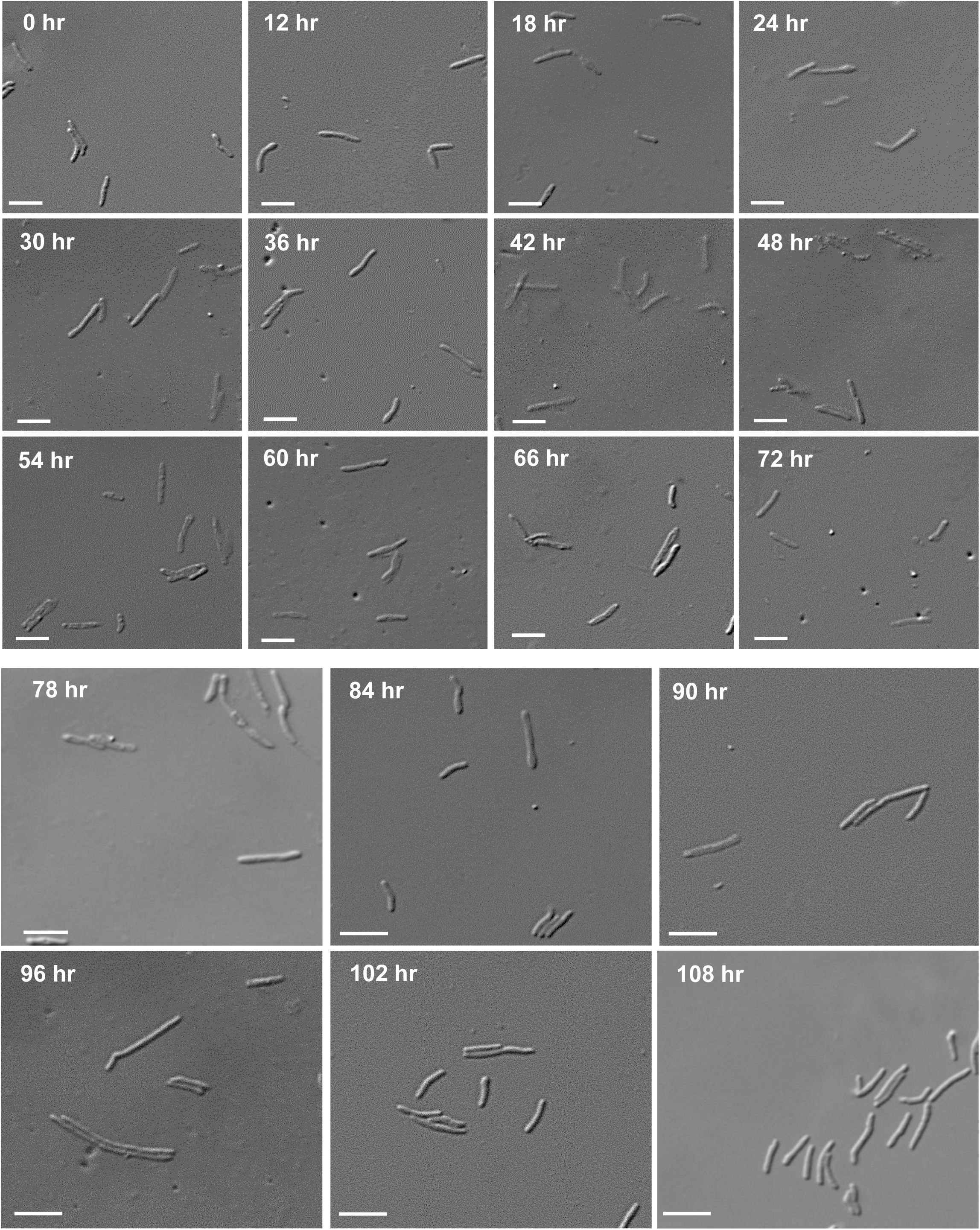
DIC images of the *Msm* cells at different time points during rifampin exposure. The cells were taken at every 6 hr interval from the rifampin-exposed population to determine the cell-lengths. DIC images of the cells during the killing phase (0 hr to 36 hr), persistence phase (36 hr to 54 hr), and regrowth phase (54 hr to 108 hr). Scale bar, 5 μm.

**Fig S6.**
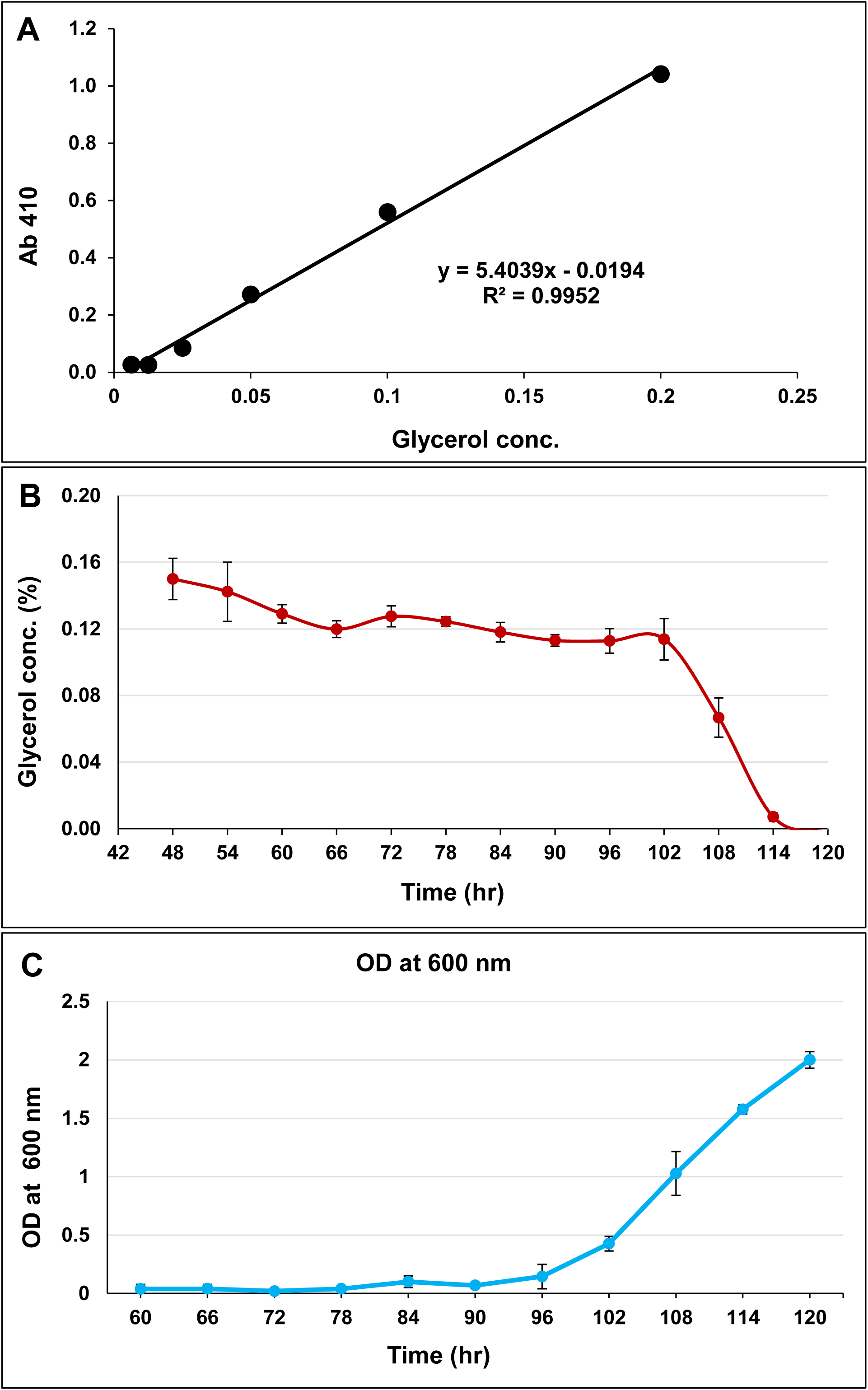
Estimation of glycerol in the *Msm* cultures during the regrowth phase. **(A)** Standard curve for glycerol estimation with different concentrations of glycerol. **(B)** The concentration of glycerol present in the culture at different time points during the exposure of the cells to rifampin. **(C)** Growth of the cells measured by observing the cell density at OD 600_nm_ at different time points.

**Fig S7.**
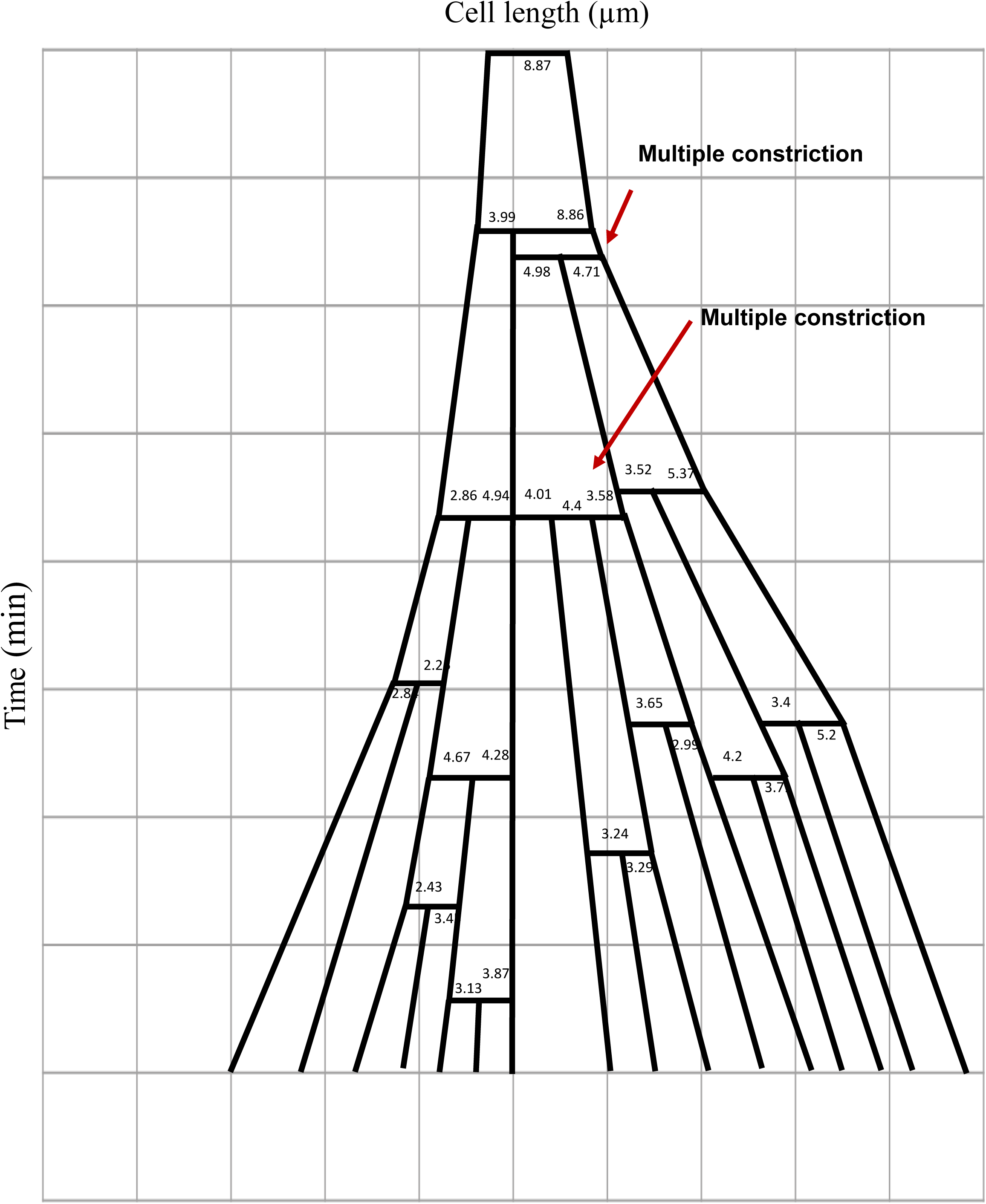
The lineage of the formation of *M. smegmatis* sister-daughter cells stressed with 5 µg/ml rifampin for 12 hrs. The growth and division lineage was traced from the images of time-lapse microscopy shown in the Video S1. The zero-time point does not correlate with the birth of the starting mother cell. Red arrows indicate the occurrence of multiple constriction. The cell lengths were drawn to approximation, not to scale.

**Fig S8.**
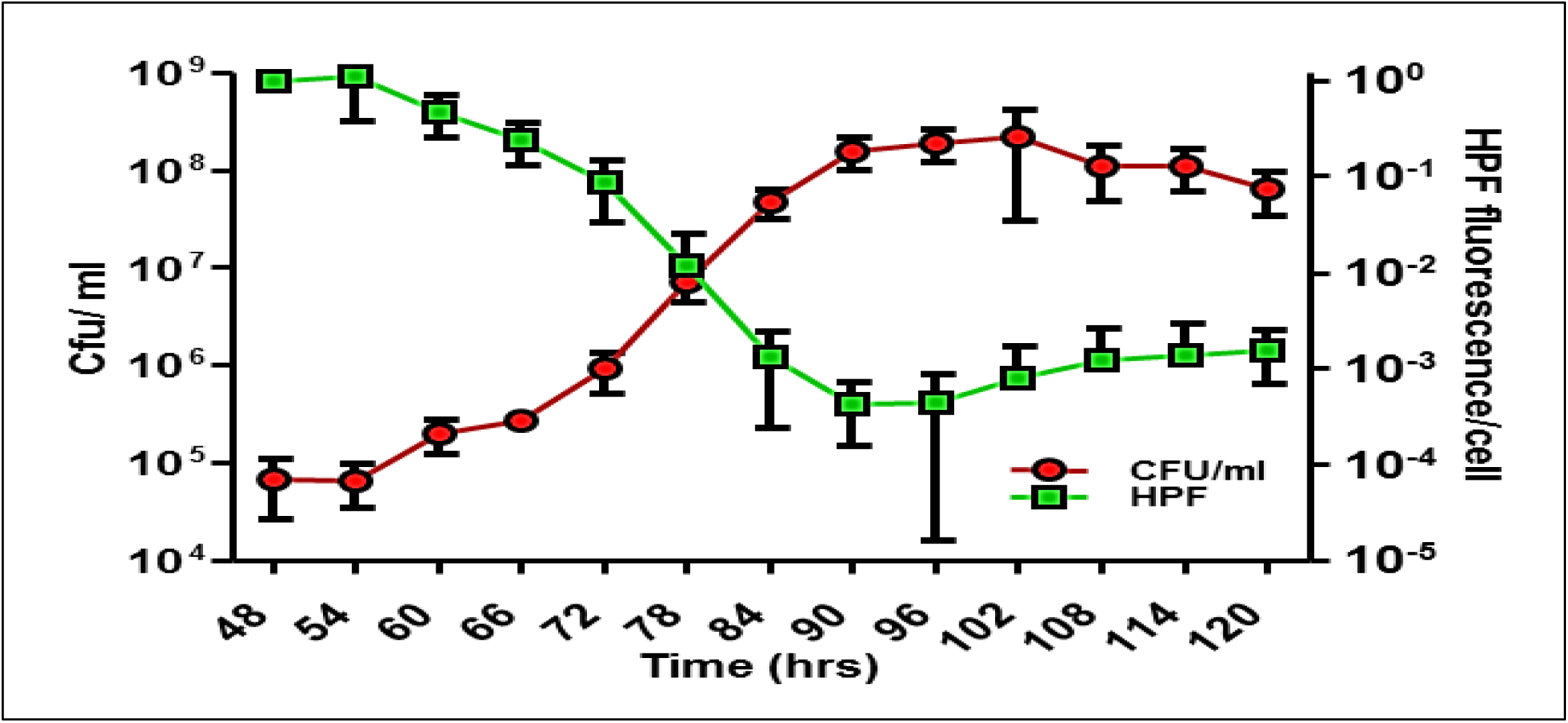
HPF fluorescence profile of the cells in the persister phase followed by regrowth phase after normalisation with their respective CFU. The hydroxyl radical levels were high during the persister phase and they declined when the cells have entered regrowth phase after acquiring RRDR mutation and becoming free from the antibiotic stress.

**Fig S9.**
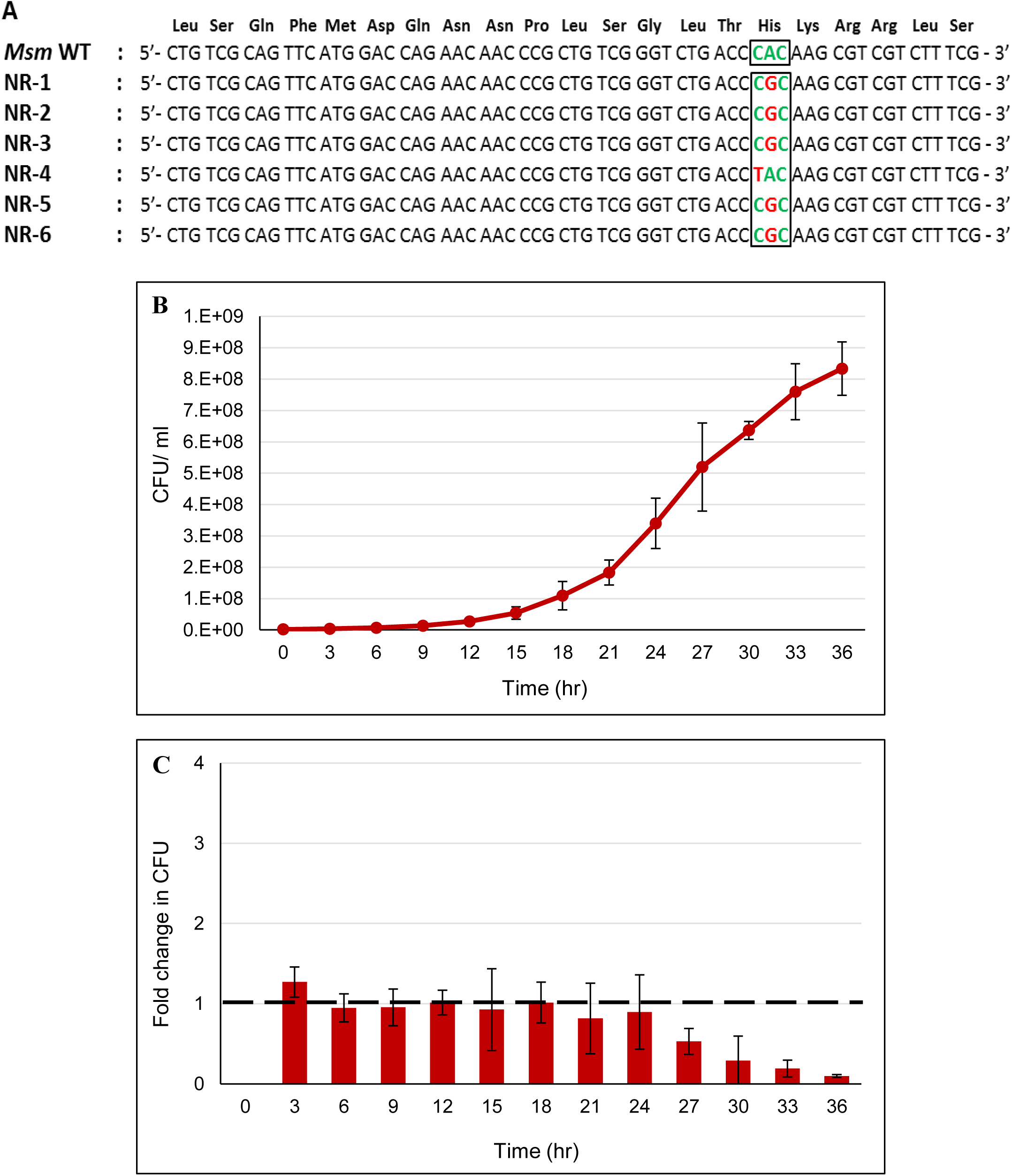
Growth, division and fold-change in the cfu of the rifampin-resistant mutants that were directly selected from the MLP culture. **(A)** RRDR mutations in the six rifampin-resistant mutants selected from the MLP culture. **(B)** Growth curve of one of the six *Msm* rifampin-resistant mutants when cultured in the presence of 25 µg/ml rifampin. **(C)** Fold-change in the cfu of the rifampin-resistant mutant grown in **(B)**.

**Table S1:**
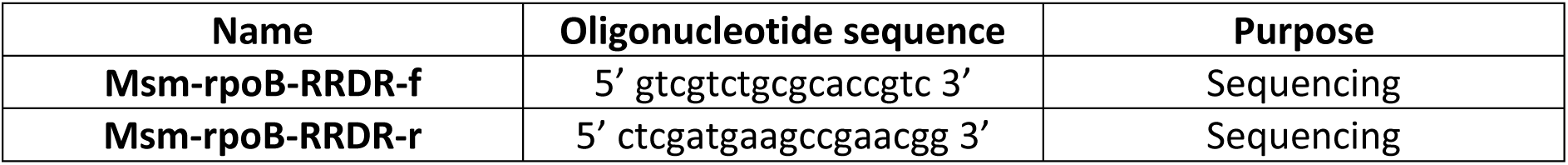
Oligonucleotides used in the study.

